# Cerebellar grey matter volume is associated with cognitive function and psychopathology in adolescence

**DOI:** 10.1101/288134

**Authors:** Torgeir Moberget, Dag Alnæs, Tobias Kaufmann, Nhat Trung Doan, Aldo Córdova-Palomera, Linn Bonaventure Norbom, Jaroslav Rokicki, Dennis van der Meer, Ole A. Andreassen, Lars T. Westlye

## Abstract

**Background:** Accumulating evidence supports cerebellar involvement in mental disorders such as schizophrenia, bipolar disorder, depression, anxiety disorders and attention-deficit hyperactivity disorder. However, little is known about the cerebellum in developmental stages of these disorders. In particular, whether cerebellar morphology is associated with early expression of specific symptom domains remains unclear.

**Methods:** We used machine learning to test whether cerebellar morphometric features could robustly predict general cognitive function and psychiatric symptoms in a large and well-characterized developmental community sample centered on adolescence (the Philadelphia Neurodevelopmental Cohort, N=1401, age-range: 8 - 23).

**Results:** Cerebellar morphology was associated with both general cognitive function and general psychopathology (mean correlations between predicted and observed values: *r* = .20 and *r* = .13; *p*-values < .0009). Analyses of specific symptom domains revealed significant associations with rates of norm-violating behavior (*r* = .17; *p* < .0009), as well as psychosis (*r* = .12; p < .0009) and anxiety (*r* = .09; *p* =.0117) symptoms. In contrast, we observed no associations with attention deficits, depressive, manic or obsessive-compulsive symptoms. Crucially, across 52 brain-wide anatomical features, cerebellar features emerged as the most important for prediction of general psychopathology, psychotic symptoms and norm-violating behavior. Moreover, the association between cerebellar volume and psychotic symptoms, and to a lesser extent norm violating behavior, remained significant when adjusting for several potentially confounding factors.

**Conclusions:** The robust associations with psychiatric symptoms in the age range when these typically emerge highlight the cerebellum as a key brain structure in the development of severe mental disorders.

## Introduction

A growing body of research reports cerebellar involvement across a wide range of mental disorders, including schizophrenia(1), bipolar disorder(2), depression(3), anxiety disorders(4), attention-deficit hyperactivity disorder(5) and autism(6). However, while the majority of these conditions are conceptualized as neurodevelopmental disorders(7, 8), most studies investigating the role of the cerebellum in mental health research have targeted adult populations(1, 9-11). Hence, it is largely unknown whether cerebellar changes can be detected already in adolescence, when initial symptoms typically first present(8, 12, 13), or only emerge later in the disease process. Moreover, whether individual differences in cerebellar structure in adolescents are indicative of non-specific impairments such as cognitive deficits (present across a wide range of psychiatric disorders(14)) or general psychopathology (analogous to the g-factor of intelligence; (15-17)), or rather are associated with specific symptom domains(18), remains unclear. Finally, it is unknown how cerebellar associations with psychiatric symptoms in adolescence compare against such associations in other brain regions. Answering these questions will be crucial for determining the relative importance of the cerebellum during this critical period for the development of mental disorders.

Here, we used machine learning to test whether cerebellar morphometric features could robustly predict cognitive function and psychiatric symptoms in a large and well-characterized developmental community sample centered on adolescence(19, 20). Consistent with Research Domain Criteria framework(21) proposed by the National Institutes of Mental Health, we followed a diagnostically agnostic and dimensional approach(22, 23), extracting clusters of correlated symptoms from a comprehensive set of clinical assessment data using blind source separation methods(24). A similar data-driven and anatomically agnostic approach was used to decompose cerebellar grey matter maps into spatially independent components, before testing for structure-function associations using multivariate machine learning. By using 10-fold internal cross-validation of machine-learning prediction models and permutation-based statistical inference in a large community sample, we aimed to optimize the robustness and generalizability of the results(25). Further, to confirm convergence across methodological approaches, we also tested for structure-function associations at the resolution levels of cerebellar lobules and voxels, and performed traditional univariate analyses in addition to running the machine learning prediction models. We finally evaluated the specificity of any cerebellar effects by testing for structure-function associations across brain-wide regions-of-interest (ROIs), tested whether associations with specific symptom domains were independent of associations with general cognitive function(26, 27) and general psychopathology(15), and controlled for potentially confounding variables such as MRI data quality(28), parental education level(29), use of psychoactive substances(30) and psychiatric assessment strategy.

Based on the existing literature on adults, we hypothesized that cerebellar morphology would be associated with both cognitive function(26, 31, 32) and general psychopathology(15), but remained agnostic as to whether such associations would show specificity across different psychiatric symptom domains.

## Methods and materials

### Participants

The main structure-function analyses were based on data from 1401 participants (52.8% female, mean age: 15.12 years, age range: 8.2 to 23.2) included in the publicly available Philadelphia Neurodevelopmental Cohort (PNC)(19, 20)(see Supplementary Methods for inclusion criteria and demographic information). The institutional review boards of the University of Pennsylvania and the Children’s Hospital of Philadelphia approved all study procedures, and written informed consent was obtained from all participants.

### Collection and processing of cognitive and clinical measures

As reported previously(24), we included performance scores from the full PNC sample (n=6,487) on 12 computerized cognitive tests(20) and 129 questionnaire items from the PNCs GOASSESS computerized assessment battery(20), including adapted items from several different questionnaires, such as the World Health Organization Composite International Diagnostic Interview (CID; (33)), the Kiddie Schedule for Affective Disorders and Schizophrenia for School Age Children (Kiddie-SADS, (28), the Structured Interview for Prodromal Syndromes (SIPS; (34)), and the PRIME Screen Revised (PRIME; (35)).

The GOASSESS battery thus allows for a broad mapping of symptoms of anxiety, mood, behavioral, eating and psychosis spectrum disorders, with a particular focus on psychosis. For individuals below 18 years of age, we relied on information from interviews with caregivers or legal guardians (20). See Supplementary Tables 1 and 2 for individual cognitive tests/clinical items. Using the full PNC sample (n=6,487), we derived general measures of cognitive performance (gF) and psychopathology (pF) by extracting the first factor scores from principal component analyses (PCA) of all cognitive and clinical scores, respectively. Next, in order to also examine specific symptom domains, all clinical item scores (n=6,487) were submitted to independent component analysis (ICA) using ICASSO(36), decomposing them into seven independent components. For the subset of participants with MRI data (n=1401), effects of sex and age on all cognitive/clinical measures were tested using generalized additive models (GAMs) as implemented in the r-package "mgcv"(37), and a set of adjusted cognitive/clinical scores were computed by regressing out main effects of age and sex (see Supplementary Methods). All subsequent structure-function analyses were conducted using these age-and sex-adjusted scores.

### Collection and processing of MRI data

As previously described(19, 38, 39), all MPRAGE T1-weighted images were collected using the same scanner (Siemens Tim Trio 3 Tesla, Erlangen, Germany; 32 channel head coil), using the following parameters; TR 1810 ms, TE 3.51 ms, FOV 180 × 240 mm, matrix 256 × 192, 160 slices, TI 1100 ms, flip angle 9 degrees, effective voxel resolution of 0.9 x 0.9 x 1mm. All images were first processed using FreeSurfer version v5.3 (http://surfer.nmr.mgh.harvard.edu), yielding estimates of total intracranial volume (eTIV)(40), volumes of eight subcortical structures(41) and mean cortical thickness of 34 cortical regions-of-interest (ROIs) per hemisphere(42). Next, the bias-corrected images from the FreeSurfer pipeline were subjected to cerebellum-optimized voxel-based morphometry (VBM) using the SUIT-toolbox (v3.2(43, 44)), running on MATLAB 2014a. In the first step, SUIT isolates the cerebellum and brainstem, segments images into grey and white matter maps. In order to avoid voxels from the occipital cerebral cortex in the SUIT-generated cerebellar grey matter maps, we excluded all voxels that overlapped with the FreeSurfer-generated maps of cortical grey and white matter. In a second step, SUIT normalizes these (FreeSurfer-pruned) cerebellar grey matter maps to a cerebellar template using Dartel(45), ensuring superior cerebellar alignment compared with whole-brain procedures(44). Normalized cerebellar grey matter maps were modulated by the Jacobian of the transformation matrix to preserve absolute grey matter volume, and the volumes of 28 cerebellar lobules were extracted using the SUIT probabilistic atlas. Next, maps were smoothed using a 4 mm FWHM Gaussian kernel before being subjected to ICA or voxel-wise general linear models. Finally, a mask for these analyses was constructed by thresholding the mean unmodulated cerebellar grey matter map at .01 and multiplying it with the SUIT grey matter template (also thresholded at .01).

### Data-driven parcellation of cerebellar grey matter

Since cerebellar parcellations based on gross anatomical features (e.g., lobules) only partially overlap with functional maps of the cerebellum(46), we used a data-driven approach in our primary analyses. Specifically, we subjected the modulated cerebellar grey matter maps to ICA using FSL MELODIC(47), testing model orders from 5 to 20 and, pragmatically, selecting the model order yielding the maximal number of clearly bilateral components for further analysis.

In order to characterize the resulting cerebellar VBM-components, we used NeuroSynth(48) to map the full-brain functional connectivity of each components peak voxel, and decoded these full-brain connectivity maps in terms of their similarity to (i.e., spatial correlation with) meta-analytic maps generated for the 2911 terms in the NeuroSynth(48) database (see Supplementary Methods).

### Analysis of brain-behavior associations

Before inclusion in statistical models, all volumetric features were adjusted for effects of age, sex and eTIV, using GAMs to sensitively model and adjust for potentially non-linear effects of age(49-51) and eTIV(52, 53) (see Supplementary Methods).

In our primary analyses, we tested whether subject weights on cerebellar independent components could predict cognitive and clinical scores, by using shrinkage linear regression(54) (implemented in the R-package ‘care’) with 10-fold internal cross-validation (i.e., based on iteratively using 90% of the sample to predict the remaining 10%), repeated 10,000 times on randomly partitioned data. Model performance was evaluated by computing the Pearson correlation coefficient between predicted and observed cognitive/clinical scores (taking the mean across iterations as our point estimate). Statistical significance was determined by comparing these point estimates to empirical null distributions of correlation coefficients under the null hypothesis (computed by running the models 10,000 times on randomly permuted clinical/cognitive scores). Results were considered significant at p < .05 (one-tailed), Bonferroni-adjusted for the 9 tested associations. In order to determine the relative importance of the anatomical features included in each prediction model, we computed the squared correlation-adjusted marginal co-relation (CAR) scores(55) for each iteration, yielding distributions of 10*10,000 CAR^2^ estimates.

To complement these multivariate prediction models, we performed a set of univariate analyses, correlating the (age- and sex-adjusted) subject weights on each cognitive/clinical component with the (eTIV-age- and sex-adjusted) anatomical subject weights (see Supplementary Methods).

In order to facilitate comparison with previously published research, we also report results from prediction models and correlation analyses using volumetric estimates of 28 cerebellar lobules as features as well as from general linear models performed at the voxel level. The voxel-wise analyses tested for effects of cognitive/clinical scores while controlling for effects of sex, age, and eTIV using FSLs randomise(56) with 10,000 permutations per contrast.

Next, to allow for a comparison of cerebellar and cerebral structure-function associations, all prediction models were also performed on volumetric estimates of eight bilateral subcortical structures, and estimates of cortical thickness from 34 bilateral ROI based the Desikan-Killany atlas in FreeSurfer, respectively (See Supplementary Figure 2). We chose thickness as our cortical feature of interest, due to its generally stronger and more consistent associations with psychopathology than surface area(57). All anatomical indices were adjusted for effects of age and sex (and eTIV for volumetric indices), as described above. Finally, in order to directly compare relative feature importance across all anatomical measures, prediction models were also fitted using z-normalized versions of all morphometric features.

**Figure 2:**
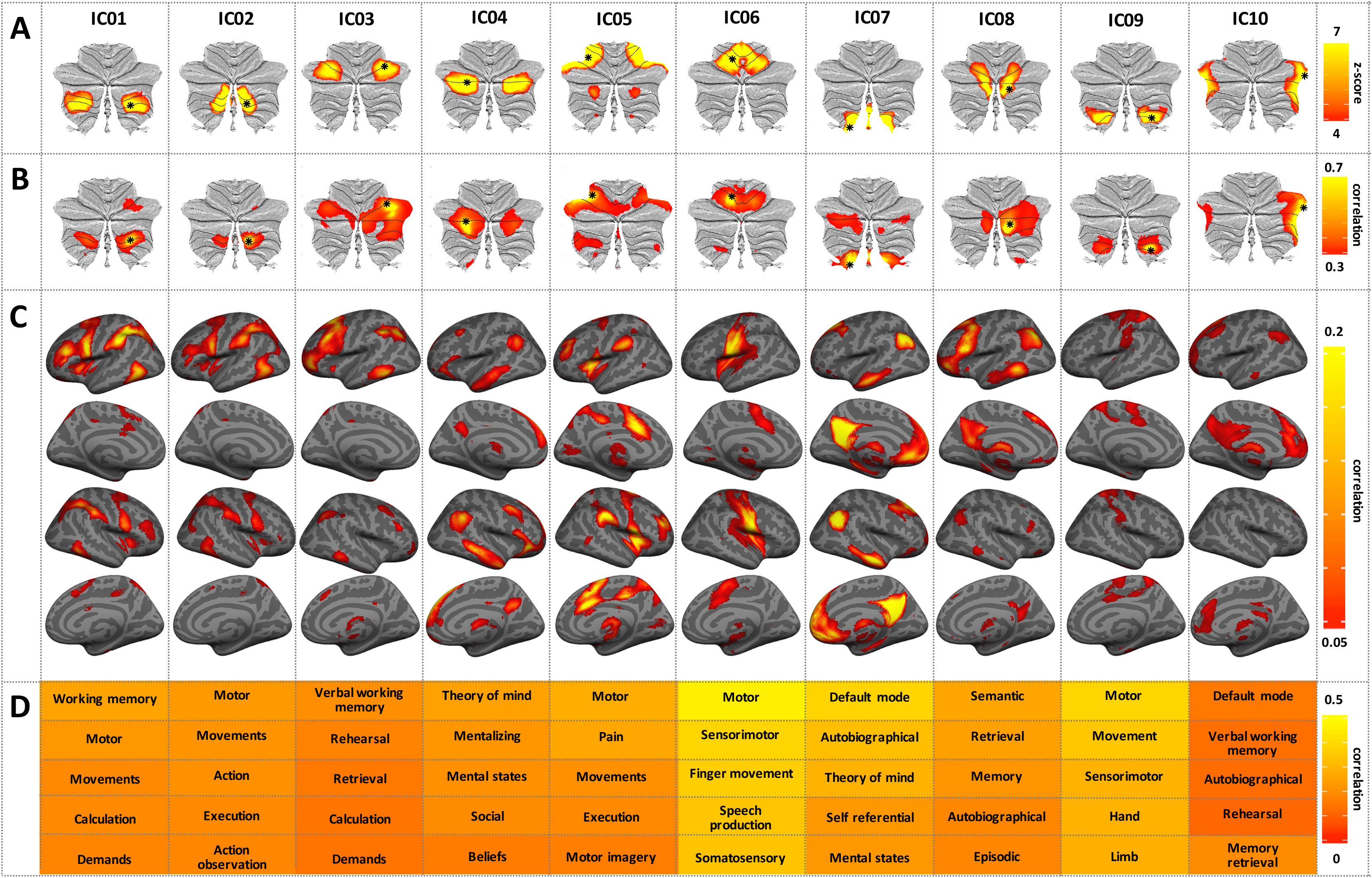
Cerebellar anatomical indices. **a:** The ten independent components resulting from data-driven decomposition of cerebellar grey matter maps projected onto flat-maps of the cerebellar cortex(91). Asterisks denote the peak voxel for each component. **b-c:** Cerebellar and cerebro-cortical functional connectivity maps (determined using NeuroSynth(46, 92)) for each of the peak voxels shown in **a**. **d:** Top 5 functional terms associated with each of the full-brain cerebellar connectivity maps shown in **b** and **c**.

Effects of general cognitive function and general psychopathology, as well as potentially confounding variables such as MRI data quality, parental education, use of psychoactive substances and psychiatric assessment strategy (interviews with caregivers or self-report) were examined by running a set of univariate control analyses on subsets of subjects (n = 369-1401) with available information (see Supplementary Methods).

## Results

### Cognitive function and clinical symptoms

Results from the PCA and ICA decompositions of clinical item scores are shown in Figure 1a. As reported previously(24), the ICA yielded seven components, primarily reflecting symptoms of attention deficit hyperactivity disorder (ADHD), various anxiety disorders (Anxiety), norm violating behavior/conduct problems (Conduct), psychotic symptoms (Psychosis), depression (Depression), mania (Mania) and obsessive-compulsive disorder (OCD). See Supplementary Table 2 for a list of all clinical items and Supplementary Figure 3 for item-specific numerical PCA and ICA weights. Effects of age and sex on all cognitive and clinical summary scores are displayed in Figure 1b and Supplementary Table 3. In brief, general cognitive function (gF) showed the expected strong positive association with age, with slightly higher mean scores in males than in females. General psychopathology also increased over the sampled age span, but did not differ between males and females. All clinical scores varied as a function of age. Specifically, ADHD scores decreased with increasing age, whereas various increasing trends were observed for all other clinical components. Largely in line with population-based estimates(8, 58, 59), males scored higher on components reflecting ADHD, conduct problems, psychosis and mania, while females had higher scores on components reflecting various anxiety disorders and OCD. No significant sex differences were observed for depression. As shown in the lower triangle of Figure 1c and Supplementary Table 4, age and sex-adjusted subject weights on clinical independent components were only weakly correlated with each other (*r* < .1), but showed moderate positive correlations with general psychopathology (pF; *r* ranging from .22 to .54). General cognitive function (gF) showed weak negative correlations with general psychopathology (pF), ADHD, Anxiety, Conduct and Psychosis (*r*-values ranging from -.1 to -.23).

**Figure 1:**
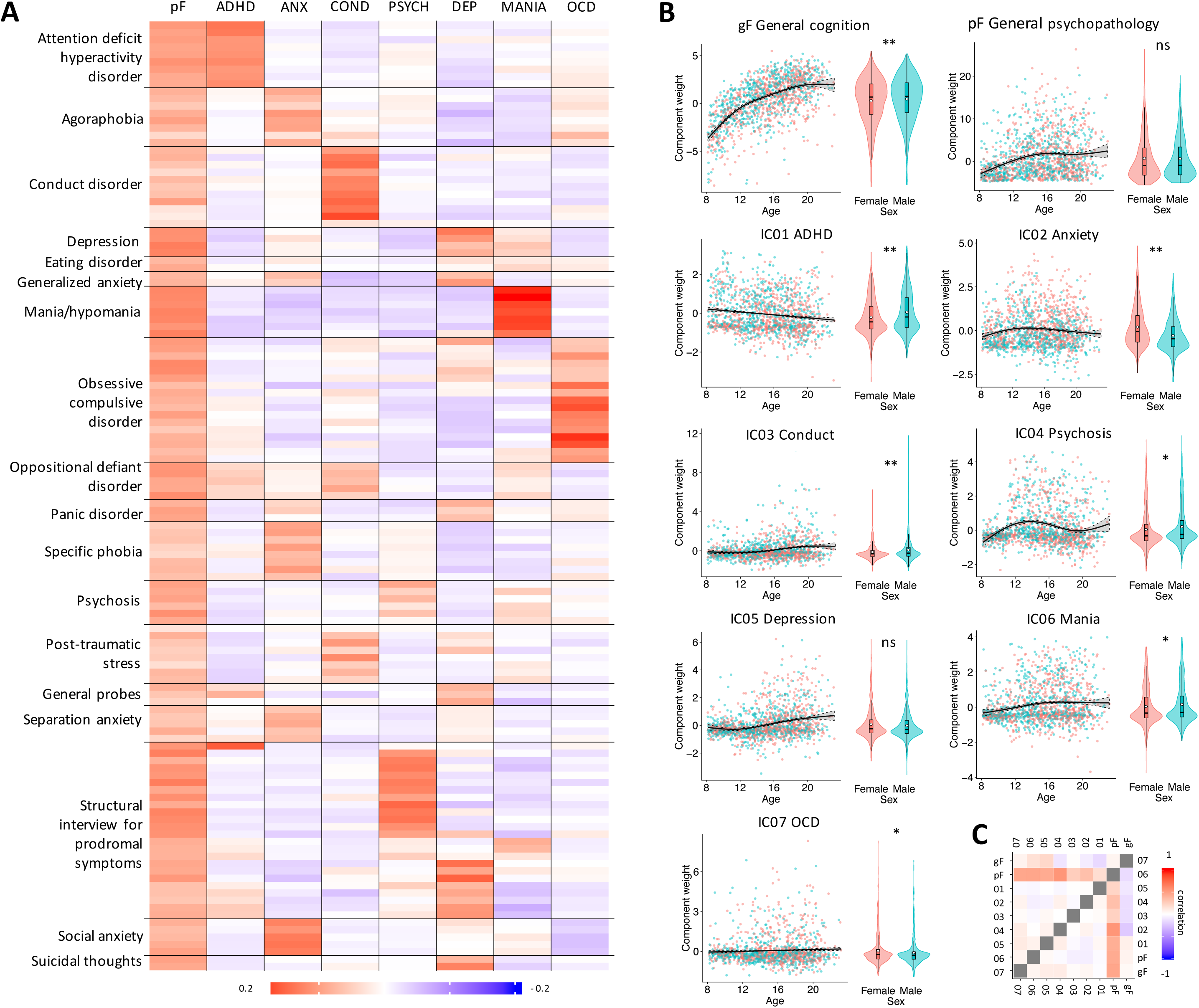
Behavioral indices. **a:** Loadings of 129 clinical items from 18 questionnaires on the general psychopathology factor (pF) and the seven clinical independent components, primarily reflecting symptoms of attention deficit hyperactivity disorder (ADHD), anxiety (ANX), norm-violating behavior/conduct disorder (COND), psychosis (PSYCH), depression (DEP), mania (MANIA) and obsessive-compulsive disorder (OCD). Clinical conditions targeted by each questionnaire are listed on the y-axis, while Supplementary Table 2 lists all 129 individual items and Supplementary Figure 3 gives numeric PCA and ICA weights for each item; **b:** Effects of age and sex on cognitive/clinical scores (asterisks denote significant sex differences; * < .05, *** <.001); **c:** correlations between all cognitive/clinical scores before (upper triangle) and after (lower triangle) correcting for effects of age and sex.

**Figure 3:**
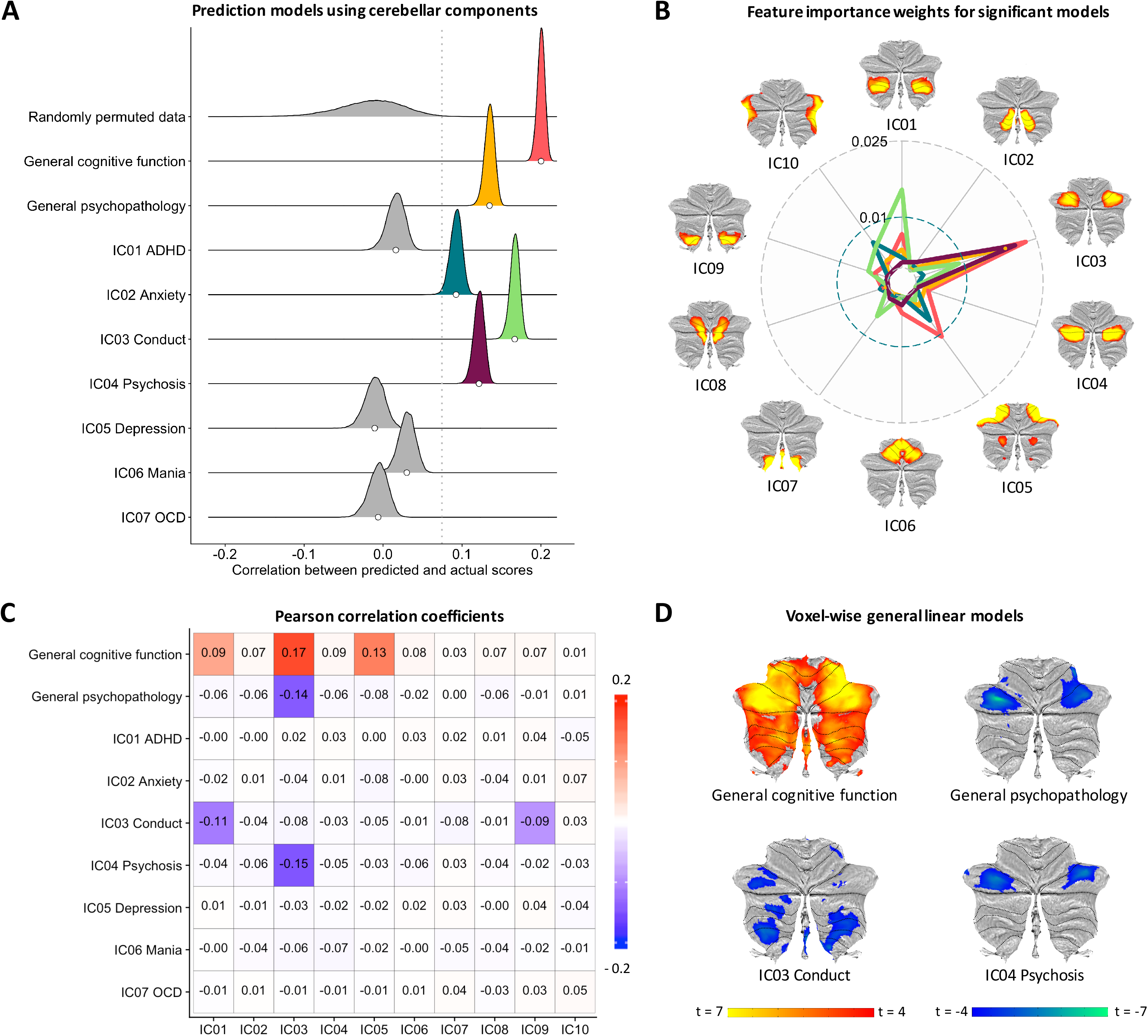
Cerebellar structure-function associations. **a:** Distributions of correlations between predicted and actual cognitive/clinical scores across 10,000 iterations of the 10-fold cross-validated model. White dots denote the mean, used as point estimates for comparison with each model’s empirical null distribution (computed by fitting the predictive models to randomly permuted cognitive/clinical data, across 10,000 iterations). For illustrative purposes we here plot the empirical null-distribution summed across all prediction models. The dotted grey line represents the one-tailed .05 threshold, Bonferroni-adjusted for 9 tests. **b:** Feature importance weights (CAR-scores) for the five significant models (color code as in a); **c:** Univariate correlations between cerebellar ICs and cognitive/clinical scores. Colored tiles mark significant associations (corrected for multiple comparisons across the matrix); **d:** T-statistics from the voxel-wise general linear models, thresholded at p < .05, two-tailed (based on 10.000 permutations).

### MRI-based morphometry

Data-driven decomposition of cerebellar grey matter maps using a model order of 10 yielded a set of symmetric bilateral components (Figure 2a), which tended to fuse using lower model orders and split into unilateral components at higher model orders (see Supplementary Figures 4-6 for results using model orders of 5, 15 and 20). We consequently chose the 10-component decomposition for all further analyses. Of note, the Neurosynth analyses revealed that voxels at the peak coordinates of each cerebellar component (marked with an asterisk in Fig.2a) showed distinct patterns of whole-brain functional connectivity (Fig 2b-c), which were associated with different functional terms in the neuroimaging literature (Fig 2d). In brief, the connectivity maps of four components (IC02, IC05, IC06 and IC09) were most closely associated with motor control, while the remaining connectivity networks showed stronger associations with various cognitive functions. See Supplementary Figures 7-10 and Supplementary Tables 5-8 for estimated effects of age, sex and eTIV on all anatomical features.

**Figure 4:**
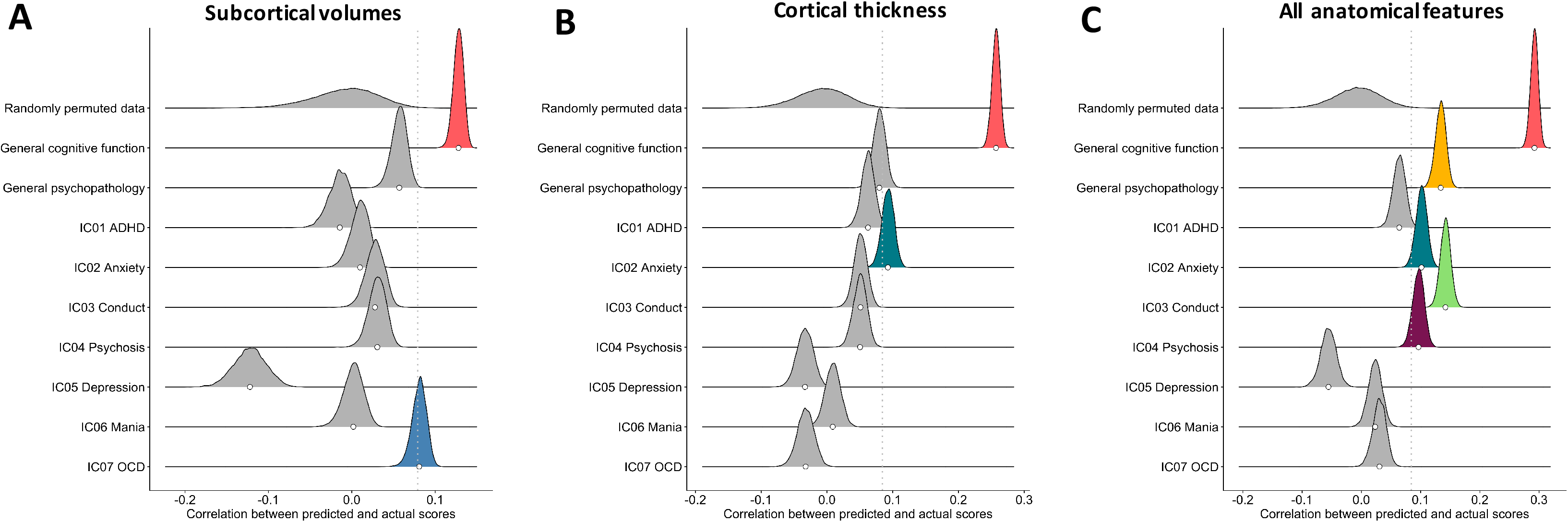
Predictive performance of cerebral models. **a:** Predictive performance of machine learning models using **a:** Subcortical volumes; **b:** Mean thickness for 34 bilateral cerebrocortical ROIs; and **c:** Z-normalized versions of all anatomical features.

### Structure-function associations

Results from the main structure-function analyses are presented in Figure 3. As hypothesized, cerebellar morphological features predicted both general cognitive function (mean correlation between observed and predicted scores: *r* = .20; *p* < .0009) and general psychopathology (*r* = .12, *p* < .0009). When using cerebellar features to predict clinical components, we observed significant results for Conduct (*r* = .16; *p* < .0009), Psychosis (*r* = .12; *p* < .0009) and Anxiety (*r* = .09; *p* = 0.0117), but not for ADHD (*r* = .01; ns), Depression (*r* = -.02; ns), Mania (*r* = .03; ns) or OCD (*r* = -.01; ns). The relative feature importance (i.e., CAR-score) for each cerebellar component used in the five significant prediction models is presented in Figure 3b. Briefly, IC03 contributed most strongly to the prediction of cognitive function (gF), general psychopathology (pF) and psychotic symptoms, whereas IC01 was the most important feature when predicting conduct problems.

This pattern was confirmed in the univariate analyses (Figure 3c). Specifically, general cognitive function (gF) was positively correlated with subject weights on IC01, IC03, and IC05, while overall psychopathology (pF) was negatively correlated with subject weights on IC03. Of the seven clinical ICs, Conduct was negatively correlated with cerebellar IC01 and IC09, while Psychosis was negatively correlated with cerebellar IC03. No other associations survived correction for multiple comparisons. Prediction models and univariate analyses using cerebellar lobular volumes yielded very similar results (see Supplementary Figure 12).

Results from the voxel-based analyses are given in Figure 3d and Supplementary Table 9. In line with the main findings, we observed anatomically widespread positive associations with general cognitive function, while general psychopathology scores were associated with a more restricted pattern of cerebellar grey matter volume reduction, encompassing bilateral lobule VI and Crus I. Psychotic symptoms were associated with a largely overlapping pattern, while conduct problems were associated with a partially overlapping region in left Crus I, as well as additional clusters in more inferior and midline regions. Anxiety was negatively associated with a small cluster in left lobule VI (11 voxels, not shown). No other clinical component yielded significant voxel-wise results.

### Prediction models using cerebral anatomical features

Figures 4 a-c present the performance of prediction models using volumetric estimates of 8 bilateral subcortical structures, cortical thickness estimates from 34 bilateral cerebral ROIs and scaled versions of all anatomical measures, respectively (see Supplementary Figures 13-14 for CAR-scores). In brief, the subcortical model performed worse than the cerebellar model, with a notable exception for OCD (*r* = .08; *p* = .0423), where pallidum volume emerged as the most important feature. The cortical thickness model performed better than the cerebellar model for general cognitive function (*r* = .26; *p* < .0009) and yielded comparable results for Anxiety (*r* = .09; p = .0225), but performed worse than the cerebellar model in predicting general psychopathology, Conduct and Psychosis (all *r*s < .0.08; all *p*s => .072). Models using all anatomical features significantly predicted general cognitive function (*r* = .29; *p* < .0009), general psychopathology (*r* = .13; *p* < .0009), Anxiety (*r* =.10; *p* = .0153), Conduct (*r* =.14; *p* < .0009) and Psychosis (*r* =.10; *p* = .0162). Univariate analyses yielded similar results (see Supplementary Figures 15-16).

Figure 5 gives the feature importance weights for significant models using all anatomical features. Of note, cerebellar features emerged as the most important in several of these models, especially general psychopathology, Conduct and Psychosis.

**Figure 5:**
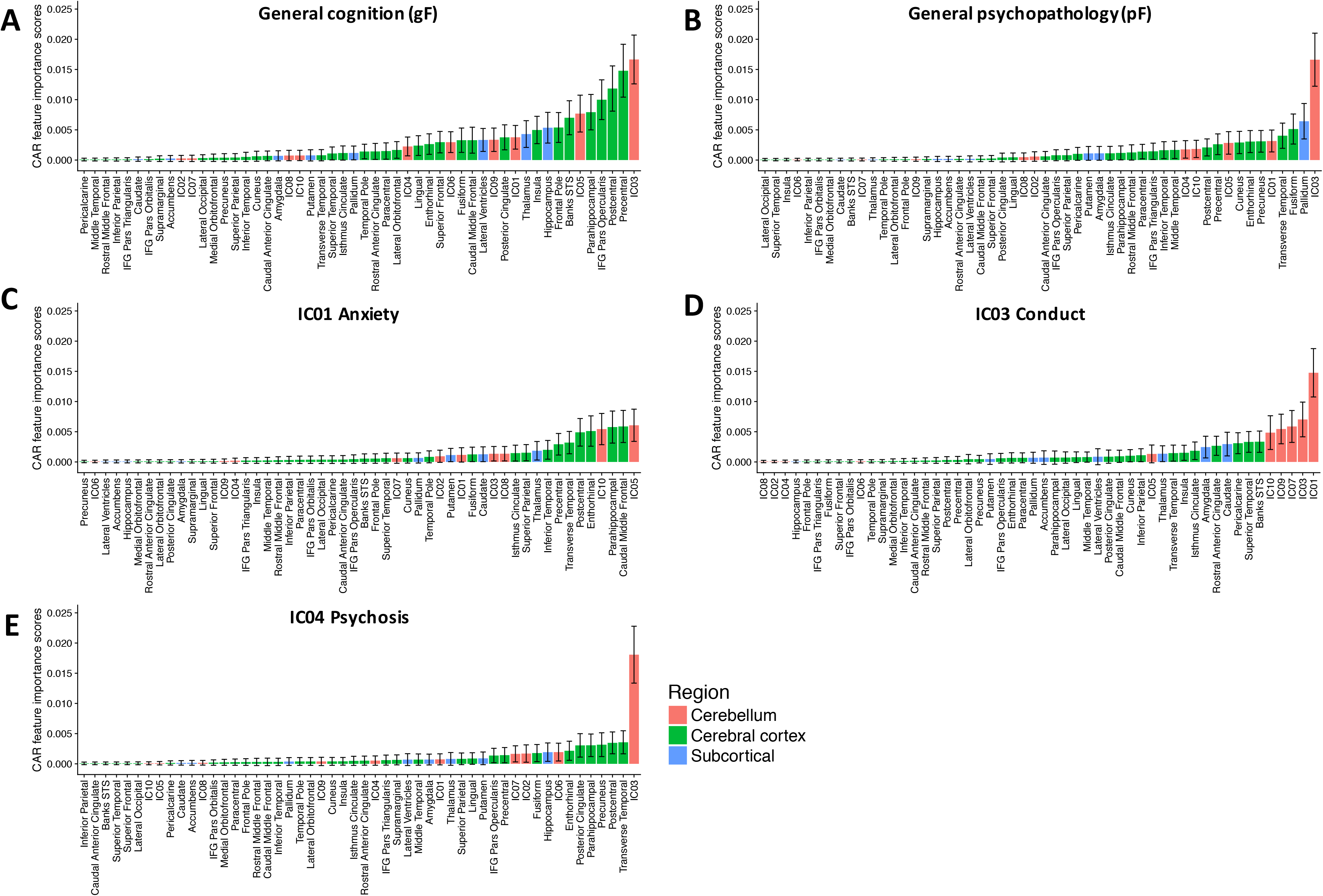
Relative feature importance across brain regions. Feature importance weights (CAR-scores) for the five significant prediction models using all anatomical features. CAR-scores were computed for each of 10,000 iterations of the model on randomly 10-fold partitioned data, yielding 100,000 estimates for each model. Colors indicate the general anatomical classification of each feature, while error bars denote the 2.5th and 97.5th percentiles of these CAR-score distributions.

### Control analyses

See Supplementary Results and Supplementary Figure 17 for detailed results. In brief, the negative correlation between cerebellar IC03 and psychotic symptoms remained significant when controlling for general cognitive function, general psychopathology, MRI data quality, parental education level, as well as in the subsets of participants with no evidence of substance abuse or assessed using only collateral information from caregivers (all corrected *p-*values < .05). The negative correlation between cerebellar IC01 and conduct problems was no longer significant when controlling for parental education level or substance abuse.

## Discussion

The current machine learning approach utilizing 10-fold internal cross-validation in a large developmental MRI sample yielded three main findings. First cerebellar morphological features could significantly predict both general cognitive function and general psychopathology in adolescence. Second, the analyses of independent components based on clinical symptom scores revealed a pattern of diagnostic specificity, in that significant results were observed for psychosis symptoms and rates of norm violating behavior (i.e., conduct problems) and to a lesser extent anxiety, whereas symptoms of ADHD, depression, mania and OCD were unrelated to cerebellar morphology. These patterns also showed anatomical specificity, with volume reductions in bilateral lobules VI/Crus I most strongly related to psychosis symptoms and volume reductions in more inferior cerebellar regions (lobules VIIb and VIII) most highly correlated with norm-violating behavior. Third, associations with psychotic symptoms and norm-violating behavior were stronger for the cerebellum than for subcortical volumes or regional cortical thickness. Together, these findings provide compelling evidence for an association between cerebellar structure and the early expression of core phenotypes of severe mental illness.

The associations with general cognitive function and general psychopathology were expected based on previous research in adults(26, 31, 32), and add to the growing database supporting a cerebellar role in cognition and affect(60). Of note, the voxelwise analyses (Figure 3d) revealed an anatomically widespread pattern for general cognitive function, while general psychopathology showed a more restricted pattern, largely overlapping with that seen for the psychosis domain. One possible explanation for this overlap is that psychosis symptoms lie at the very extreme end of a putative continuum going from more benign to more severe psychopathology (16, 17). Thus, participants with high scores on the Psychosis component would also be expected to express high symptom levels more generally. In line with this notion, Caspi et al. observed a very high correlation (.997) between a Thought Disorder factor (reflecting symptoms associated with psychotic disorders) and their General Psychopathology factor (reflecting overall psychopathology) calculated based on extensive structured interviews in a large longitudinal sample(17). Consistent with their results, across the seven clinical components we also observe the highest correlation with General Psychopathology for Psychosis (IC04). Of note, measures of general psychopathology have recently been associated with a range of brain phenotypes(61-63), including white matter integrity and grey matter volume of the cerebellum(15).

The majority of existing structural MRI-studies on psychosis have focused on cerebral structures(64), but our findings on psychotic symptoms are nonetheless in general agreement with an emerging body of research. For instance, we have recently shown that cerebellar volume reductions is one of the strongest and most consistent morphological alterations in a large multi-site sample of schizophrenia patients (N = 983) and healthy controls (N = 1349)(1). Of note, both in our previous patient study(1) and in the current study of premorbid symptoms, the strongest effects of the psychosis domain converged on cerebellar regions that show functional connectivity with the frontoparietal cerebral network, which has been strongly implicated in cognitive control processes(65). Indeed, previous functional MRI studies of working memory using the PNC sample found that activation of cerebellar Crus I was associated task performance (66), and that reduced activation in this region was associated with overall level of psychopathology (67). Structural alterations in this cerebellar region also emerged as one of the strongest predictors of transition to psychosis in a recent study of high-risk populations(68). Considered together, these findings provide converging evidence for cerebellar Crus I as a key node involved in both high-level cognition and severe psychopathology. More broadly, functional neuroimaging studies consistently report reduced cerebello-cerebral connectivity in schizophrenia patients(69, 70) and high-risk groups(71, 72), while behavioral studies find impaired cerebellar learning in both patients with schizophrenia(73-75) and their first-degree relatives(76).

Of note, our findings differ in some respects from a previous study of structural brain alterations in a partially overlapping sample of psychosis spectrum youth^(38)^, which reported the strongest group effects in medial temporal, posterior cingulate and frontal regions. We highlight two possible sources of these discrepancies. First, only the current study employed analysis pipelines optimized for both the cerebellum(77) and the cerebrum(78). Second, whereas the previous study employed an extreme group design(38), we tested parametric associations across the full phenotypic range.

The associations between cerebellar volume and rates of norm-violating behavior are consistent with some recent reports of altered cerebellar white matter microstructure(79) and functional activation(80) in conduct disorder. However, since our control analyses suggested that these associations might be partially confounded by parental education level and substance abuse, they should be interpreted with caution.

While the current results do not allow inferences regarding the pathophysiological mechanisms underlying these changes in cerebellar morphology, we observe that genes associated with schizophrenia have been shown to be highly expressed in the human cerebellum(81), suggesting a direct or indirect genetic impact. Further, cerebellar volume, like hippocampal volume(82), has been shown to be very sensitive to stress hormone exposure. While this effect is especially strong during infancy(83) it has also been observed in adults with very high levels of circulating corticosteroids due to Cushing’s disease(84). These latter observations may provide a possible link, to be tested in future research, between our findings and the well-documented role of stressful life events in the development of psychopathology(85).

Strikingly, associations with psychotic symptoms and norm-violating behavior were stronger for the cerebellum than for subcortical volumes or regional cortical thickness. Since this pattern was not observed across all examined phenotypes (e.g., cortical thickness was the best predictor of general cognitive function, while subcortical volumes showed stronger associations with OCD symptoms), we believe these brain-wide comparisons reveal a crucial cerebellar involvement with respect to these specific symptom domains.

From a methodological point of view, it is worth mentioning that our data-driven cerebellar decomposition yielded components that only partially overlapped with standard anatomical parcellations. In particular, borders between several components were primarily organized along the medial-to-lateral dimension, and one component could span parts of several lobules. Together with results from recent fMRI-studies (86, 87), these results suggest that traditional cerebellar subdivisions do not optimally capture either the inter-subject structural variability or the functional heterogeneity of the cerebellum.

Notable strengths of the current study include the use of a large sample and internal cross-validation methods, which should reduce the risk of overfitting and thus ensure more generalizable effect estimates(25). Its main limitation is the cross-sectional design, which prevents direct tests of causal relationships. We observe, however, that results from previous studies using smaller longitudinal samples do suggest that cerebellar structure and function can predict later progression of psychotic symptoms(72) or conversion to frank psychosis(68). A second limitation is that only a subset of participants had information on parental education and substance abuse, resulting in a reduced sample size for some of the control analyses. Third, although a previous study has found that cerebello-cerebral connectivity patterns are largely developed and similar to those seen in adults by middle childhood (88), the extent to which results from a young adult sample(46) can be generalized to the current adolescent sample remains unknown. Finally, although the reported structure-function associations were robust and highly significant, cerebellar morphology explained only a limited part of the variance in clinical scores. While not surprising, given the multiple factors that influence the expression of psychiatric symptoms(85, 89, 90), this caveat must be kept in mind when interpreting the results.

In conclusion, our findings highlight the cerebellum as a key brain structure for understanding the development of mental disorders, in particular psychosis.

## Supporting information

Supplementary Methods and Results

## Acknowledgements

This study has received funding from the European Commission’s 7th Framework Programme (#602450, IMAGEMEND), Research Council of Norway (213837, 223273, 229129, 204966/F20, 249795, and 251134), the South-Eastern Norway Regional Health Authority (2013-123, 2014-097, 2015-073, 2016-083 and 2017112) and KG Jebsen Foundation. The Philadelphia Neurodevelopment Cohort sample is a publicly available data set. Support for the collection of the data sets was provided by grant RC2MH089983 awarded to Raquel Gur, MD, PhD, and RC2MH089924 awarded to Hakon Hakonarson, MD, PhD. All participants were recruited through the Center for Applied Genomics at The Children’s Hospital in Philadelphia. No other disclosures are reported. A preprint of this paper has been posted on bioRxiv (doi: https://doi.org/10.1101/288134).

