## Supplementary Methods and Results for "Cerebellar grey matter volume is associated with cognitive function and psychopathology in adolescence"

*Participants, inclusion criteria and quality control procedures*: Access to the PNC data was obtained with permission #8642 (project title: Neurodevelopmental brain networks: Integrating multimodal imaging, cognition and genetics). Phenotypic data were available from 6487 participants, while MRI data were available from 1601 participants. Data from four subjects suffered data corruption during download, while 76 participants with severe or major physical medical conditions requiring standing medications and monitoring (as assessed by trained personnel in the PNC study team(1, 2)) and 72 participants with missing data on medical status were removed from the sample prior to analyses. In addition, one participant with MRI data was excluded due to missing phenotypic data. For the remaining 1448 participants, we implemented a flagging procedure based on robust PCA(3) for detecting signal to noise-, and segmentation outliers based on the FreeSurfer output. Flagged datasets were carefully inspected, with minor edits performed when necessary. Scans from 47 participants were rejected due to poor MRI quality based on visual inspection. For an additional quantitative assessment of MRI quality, we calculated the Euler numbers (one for each hemisphere) for each dataset and computed the sum of these two Euler numbers as an index of overall MRI quality(4). Supplementary Figure 1 displays the distributions of these summed Euler numbers for included and excluded datasets, respectively. As expected, mean Euler number differed significantly between included and excluded datasets and the distributions showed only moderate overlap. We thus believe these quantitative results corroborate our original quality control procedure. Euler numbers for included datasets were also included as covariates in control analyses (see below), testing for potentially confounding effects of MRI data quality(4). The final sample (N= 1401) ranged between 8.2 and 23.2 years of age (mean age: 15.12, SD: 3.62), and was 52.8% female. Mean age was slightly higher (t=2.7, p=.006868 for females (15.4; SD: 3.6)) than males (14.8; SD: 3.62).

*Computation of cognitive and clinical summary scores:* Supplementary Table 1 provides an overview of all cognitive test scores used to compute the index of cognitive function (gF). The gF score was computed by submitting cognitive test score from 6487 participants to a principal component analysis and extracting subject weights on the first principal component (explaining 30.7% of the variance in the total sample). The first factor weights on each cognitive measure are given in Supplementary Table 1.

Supplementary Table 2 lists all 129 items used to compute both the general index of psychopathology (pF) and the clinical independent component scores. As described previously(5), 1,627 participants had missing values on one or several clinical items. For all but two participants which had missing values on all 129 clinical items, the missing values were replaced with the nearest-neighbor value based on Euclidean distance. The percentage of missing values for the 129 items ranged from 0-7%.

In brief, analogous to the notion of a positive manifold in the cognitive domain (frequently termed the g-factor), a general psychopathology factor (or p-factor) has been proposed as a parsimonious explanation for the considerable correlations between different symptoms of psychopathology, as well as the common comorbidity and genetic overlaps across psychiatric diagnostic categories (for a recent review, see(6)). Thus, the general psychopathology factor (pF) reflects the shared variance across symptom domains, while independent components (by definition) reflects independent sources of variance in the same dataset. By reporting results analyzed from two such alternative nosological perspectives, we believe that our reported findings will have the maximal future impact, regardless of whether "lumping" (general psychopathology) or "splitting" (independent components) will eventually turn out to be the most fruitful approach towards a deeper understanding of psychopathology.

The pF score was computed by submitting data from the full set of PNC participants (N = 6487) to a principal component analysis and extracting subject weights on the first principal component (explaining 12.9% of the variance). The first principal component weights on each clinical item are displayed in Figure 1, with numerical information given in Supplementary Figure 3. As described previously(5), the ICA model order for clinical score decomposition was chosen after testing several different model orders ranging from 3-15, for each of which 100 permutations were run, and based on the observed independence and reliability of the resulting components. The independent component weights on each clinical item are displayed in Figure 1, with numerical information given in Supplementary Figure 3. Note that all clinical items (traditionally associated with distinct diagnostic categories) had positive and relatively uniform weights on pF (range from .02 to .13). In contrast, weights on independent components were more specific to a smaller set of symptoms, and showed a wider range which also included negative weights on some symptoms (range from -.07 to .19).

Effects of gender and age on cognitive/clinical sores were tested by fitting generalized additive models (GAMs) to account for potentially non-linear effects of age. We used GAM as implemented in the R-package “mgcv”, with age modeled using cubic splines with 5 knots the level of smoothness automatically selected using the restricted maximum likelihood method (“REML”). For the full model, we report the percent variance explained, for parametric terms we report t- and p-values, while for smooth age-term, we report F-values and approximate p-values.

Age- and gender-adjusted cognitive/clinical scores were computed by reconstructing data from the intercept and residuals of these GAM-models (i.e., omitting the age- and gender-coefficients).

*Independent component analysis (ICA) of cerebellar grey matter maps:*

ICA of the modulated cerebellar grey matter maps was performed using FSL MELODIC with standard settings. A binary mask was constructed by thresholding the mean unmodulated cerebellar grey matter map at .01 and multiplying it with the probabilistic cerebellar grey matter map from the SUIT template (also thresholded at .01). We initially tested model orders ranging from 5 to 20 (in steps of 1), and decided on a model order of 10 for the main analyses, since this produced a concise set of largely bilateral components which tended to fuse in to larger bilateral components at lower model orders and fragment into unilateral components at higher model orders (See Supplementary Figures 4-6 for examples of ICA-decompositions using alternative model orders of 5, 15 and 20).

The 10 independent components together accounted for 60.03% of the total variance in the modulated grey matter maps used as input to the analysis, with each component explaining between between 4.07% and 7.65% of the total variance (and between 8.31% and 12.74 % of the explained variance). In comparison, the 5-, 15- and 20-component models accounted for respectively 48.00, 65.72 and 69.73% of the total variance. Correlations between the 10 cerebellar ICs before and after adjusting for effects of sex, age and estimated total intracranial volume are given in Supplementary Figure 11.

For functional characterization of the resulting cerebellar grey matter components, we used results from a large (N=1000) resting state fMRI functional connectivity study(7) (implemented in NeuroSynth(8)) to map the full-brain functional connectivity of the peak voxel of each component, and plotted both cerebellar maps (thresholded at r = .3) and cerebrocortical maps (thresholded at r = .05) for illustration (Figure 2B and C). We next decoded these full-brain connectivity maps in terms of their similarity to (i.e., spatial correlation with) meta-analytic maps generated for the 2911 terms in the NeuroSynth(8) database. We report the top five functional terms (Fig 2D), i.e. a pruned version of Neurosynth output after exclusion of all terms related to brain anatomy, methodology, etc. Spatial correlations between whole-brain connectivity maps and the meta-analytic maps for the reported terms in Figure 2D ranged from .108 to .457.

*ROI-wise adjustment for effects of age- gender and estimated total intracranial volume:*

Before inclusion in multivariate or univariate models, all volumetric anatomical features were adjusted for main effects of gender and eTIV as described above for clinical scores, i.e., by fitting GAM-models and reconstructing data from model parameters and residuals. As for clinical sores, effects of age on anatomical features were estimated using cubic splines with 5 knots, with the level of smoothness automatically selected using the restricted maximum likelihood method ("REML"). For all volumetric features (but not for cortical thickness), we also used GAM (with the same input parameters as for age) to estimate potentially non-linear effects of eTIV(9, 10). For each GAM- model, we report total explained variance, t, F and p-values as described for cognitive/clinical data above.

*Univariate analyses:*

In a set of univariate analyses, we computed the Pearson correlation coefficients between all (sex-, age- and eTIV-corrected) anatomical features included in each prediction model and all predicted (sex- and age-corrected) cognitive/clinical components. Statistical significance (corrected for multiple comparisons across the set of features for each model) was determined by permutation testing. Specifically, we randomly permuted the cognitive/clinical subject weights 10.000 times, computed the resulting matrices of structure-function correlations and extracted the maximal and minimal correlation coefficients from each iteration to form empirically derived null-distributions. Structure-function associations were considered significant at a corrected alpha-level of .05 (two-tailed).

*Voxel-wise analyses:*

Statistical analyses testing voxel-wise associations between cerebellar volume and age- and gender-adjusted cognitive/clinical scores, while controlling for main effects of gender, age, and estimated total intracranial volume(eTIV), were performed using FSL Randomise. Sex was modelled as two binary variables, while age, eTIV and cognitive/clinical scores were z-transformed and modelled as three continuous variables. Statistical inference was based on permutation testing (using 10.000 permutations per contrast), with voxels considered significant at a corrected alpha-level of .05 (two-tailed).

*Analyses controlling for potentially confounding factors:*

Information on general cognitive function (i.e., the gF score) and general psychopathology (i.e., the pF score), as well as the mean Euler number (a quantitative index of MRI data quality(4)) were available for all 1401 subjects, while information on maternal and paternal education level was available for 1274 subjects. In order to control for these potentially confounding variables, which previously have been shown to be associated with both psychiatric symptoms(11) and brain morphology(12-15), we performed additional univariate after adjusting both clinical and anatomical features for main effects of the respective variable (in addition to sex and age, as well as estimated total intracranial volume for volumetric features).

In order to test whether the use of two different assessment strategies for participants below and above 18 years of age (collateral informants versus self report), we performed a control analysis including only the 1035 participants below 18 years. Information on substance abuse was available for 594 participants. In the control analyses examining this potentially confounding variable, we excluded all subjects who reported having experienced one or more negative psychological or physical effects of alcohol use, or having ever tried illicit substances (n=225), leaving 369 participants in this pruned subsample.

**Supplementary Results:**

*Effects of age and sex on cognitive/clinical scores*

The main results are displayed in Figure 1B, while the total percent variance explained by the models, t- and F-values for the gender and (smooth) age terms, as well as their respective p-values are given in Supplementary Table 3.

*Relationships between cognitive/clinical scores*

The main results are displayed in Figure 1C, with numerical values given in Supplementary Table 4. Briefly, correlations between individual scores on clinical independent components were very weak (range -.072 to .097), suggesting that these components indeed reflected largely independent sources of variance in the clinical data. Moreover, all clinical independent components showed weak to moderate positive correlations with the General Psychopathology factor (pF), ranging from .222 for ADHD to .538 for Psychosis. In line with previous reports in both an overlapping(16) and an independent sample(17), we also observed a weak negative correlation between General Psychopathology and General Cognition ($r = -.147$). Among the clinical independent components, ADHD, Anxiety, Conduct and Psychosis showed weak negative correlations with General Cognition (range: -.102 to -.230), while Depression, Mania and OCD showed very weak positive correlations with General Cognition (range: .033 to .080).

*Effects of age, sex and estimated total cranial volume on brain features:*

The GAM-estimated smooth functions for age and estimated total cranial volume (eTIV) are displayed (together with raw data and gender effects) in Supplementary Figures 7-10 and Supplementary Tables 4-7. Briefly, we observed significant effects of eTIV on all volumetric features, while age- and gender-effects were more variable. Specifically, with regards to the cerebellar components, we observed significant effects of age on subject weights for IC02, IC04, IC06 and IC10. When adjusting for eTIV, IC05, IC06 and IC09 showed significantly higher subject weights in males relative to females, while IC02, IC07, IC08 and IC10 showed the opposite pattern.

*Prediction models and univariate analyses using cerebellar lobular volumes:*

Results from the prediction models using cerebellar lobules as predictive features are presented in Supplementary Figure 12A. Significant results were observed for general cognitive function (gF; $r = .19$, $p < 0.009$), general psychopathology (pF; $r =.12$, $p < 0.009$), conduct disorder (IC03: $r = .15$, $p < 0.009$) and psychosis (IC04: $r = .12$, $p < 0.009$), while the nominally significant association with anxiety did not survive multiple comparisons correction (IC02: $r = .06$, ns). Cerebellar lobular volumes did not significantly predict subclinical symptoms of ADHD (IC01: $r = .02$, ns), depression (IC02: $r = .01$, ns), mania (IC02: $r = .01$, ns), or OCD (IC02: $r = .05$, ns). Feature importance indices (CAR-scores for significant prediction models are depicted in Supplementary Figure 12B-F.

*Control analyses:* Supplementary Figure 17A and B display all univariate associations between cerebellar and clinical IC scores, when adjusting both feature sets for general cognitive function and general level of psychopathology, respectively. As expected, given the existing evidence for associations between these factors and both clinical symptoms(5, 11) and brain structure(5, 13, 15), the structure-function associations observed in our main analyses were slightly reduced in these control analyses, but remained significant. Specifically, when adjusting both clinical and anatomical features for effects of general cognitive function, these analyses yielded correlation coefficients of -.11 ($p < .001$) between psychosis symptoms and cerebellar IC03, and -.10 ($p < .01$) between conduct disorder and cerebellar IC01. When similarly adjusting for effects of general psychopathology, we observed correlation coefficients of -.08 ($p < .05$) for the association between psychosis symptoms and cerebellar IC03, and -.10 ($p < .01$) for the association between conduct disorder and cerebellar IC01.

Results from the correlation analyses run after adjusting all anatomical features for effects of mean Euler number (a quantitative index of MRI data quality) are displayed in Supplementary Figure 17C. Both the association between cerebellar IC03 and psychosis symptoms (correlation coefficient: -.15, $p < .001$) and the association between IC01 and norm violating behavior (correlation coefficient: -.11, $p < .001$) remained significant.

Results from the correlation analyses run after adjusting all clinical and anatomical features for effects of maternal and paternal education levels are displayed in Supplementary Figure 17D. The association between cerebellar IC03 and psychosis symptoms remained significant (correlation coefficient: -.14, $p < .001$), but the association between IC01 and norm violating behavior did not (correlation coefficient: -.08, ns).

Results from correlation analyses in the pruned dataset containing only participants who reported no use of illicit substances and no negative experiences with alcohol (n=389) are displayed in Supplementary Figure 17E. Of note, the associations between psychosis symptoms and cerebellar IC03 remained significant in this pruned subsample (correlation coefficient: -.17, $p < .05$), while the association between cerebellar volumes and norm violating behavior did not (correlation coefficient: -.06, ns).

Results from correlation analyses in the pruned dataset containing only participants who with information provided from collateral informants (primary caregivers or legal guardians) (n=1035) are displayed in Supplementary Figure 17F. Both the association between cerebellar IC03 and psychosis symptoms (correlation coefficient: -.15, $p < .001$) and the association between IC01 and norm violating behavior (correlation coefficient: -.12, $p < .001$) remained significant.

**Supplementary figures and tables:**

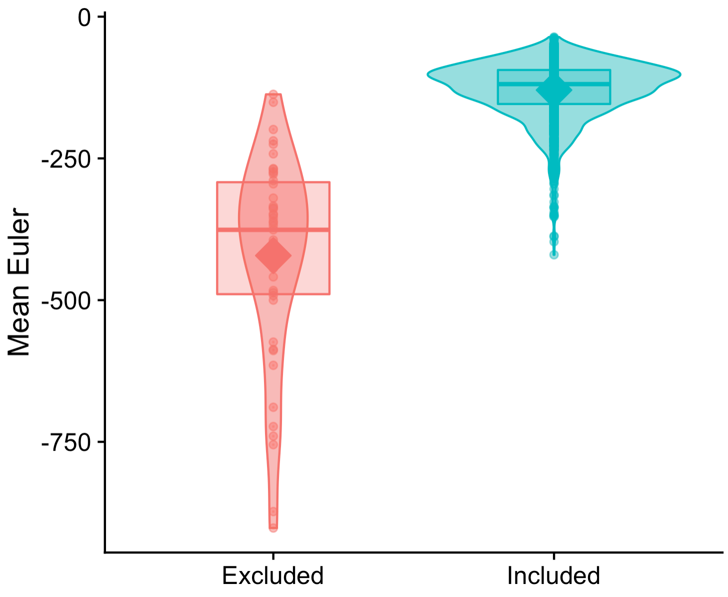


**Supplementary Figure 1:** Mean Euler numbers for included and excluded datasets. Violin plots represent the distributions, small circles individual data points, and the large diamonds group means, while boxplots show medians end interquartile ranges. Mean Euler numbers differed significantly between included and excluded datasets (p = $2^{-e4}$, based on 10.000 permutations).

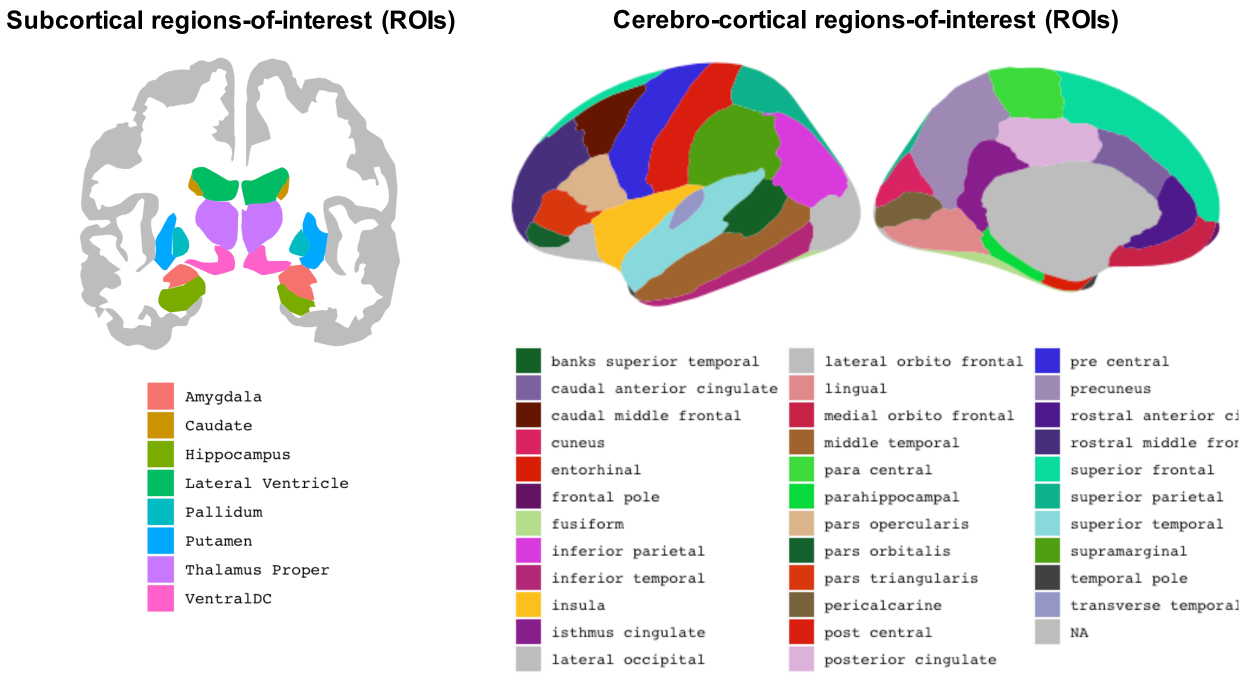


**Supplementary Figure 2:** The subcortical and cerebro-cortical regions-of-interest used in the current study. Figures have been produced using the R-package "ggseg"(18).

| | pF | ADHD | ANX | COND | PSYCH | DEP | MANIA | OCD |
|---|---|---|---|---|---|---|---|---|
| 001 | 0.11 | 0.14 | 0 | –0.03 | 0.01 | –0.02 | –0.01 | 0 |
| 002 | 0.1 | 0.13 | –0.02 | –0.02 | 0 | –0.01 | –0.02 | 0.01 |
| 003 | 0.1 | 0.11 | –0.01 | –0.02 | –0.03 | 0.02 | 0.02 | 0 |
| 004 | 0.09 | 0.11 | 0.01 | –0.02 | 0.01 | –0.02 | 0 | –0.01 |
| 005 | 0.1 | 0.11 | –0.01 | –0.03 | 0.02 | –0.01 | –0.01 | 0.01 |
| 006 | 0.11 | 0.12 | 0 | –0.01 | 0.01 | –0.03 | 0 | 0.01 |
| 007 | 0.1 | 0.11 | –0.01 | –0.02 | 0 | –0.04 | 0.02 | 0.01 |
| 008 | 0.09 | 0.11 | –0.02 | 0 | 0 | –0.03 | –0.01 | 0.02 |
| 009 | 0.09 | 0.12 | 0 | –0.01 | 0.01 | –0.04 | 0 | 0.02 |
| 010 | 0.08 | 0.01 | 0.09 | 0 | 0 | –0.01 | –0.02 | 0.02 |
| 011 | 0.07 | 0.01 | 0.07 | 0 | 0.02 | 0 | –0.04 | 0.02 |
| 012 | 0.06 | 0 | 0.05 | –0.01 | 0.02 | –0.04 | –0.02 | 0.03 |
| 013 | 0.08 | –0.01 | 0.11 | 0.02 | 0.01 | –0.06 | –0.03 | 0.02 |
| 014 | 0.1 | 0 | 0.09 | 0 | 0.02 | –0.04 | –0.03 | 0 |
| 015 | 0.08 | 0 | 0.1 | 0.01 | 0 | –0.03 | –0.03 | 0.02 |
| 016 | 0.04 | 0 | 0.03 | 0.05 | –0.02 | –0.02 | –0.04 | 0.07 |
| 017 | 0.08 | –0.01 | 0.09 | 0.02 | 0 | –0.06 | –0.03 | 0.03 |
| 018 | 0.08 | 0.01 | 0 | 0.1 | –0.01 | –0.01 | 0.02 | –0.02 |
| 019 | 0.08 | –0.03 | 0 | 0.14 | –0.04 | 0.02 | 0 | –0.03 |
| 020 | 0.06 | 0.01 | –0.01 | 0.15 | 0 | –0.02 | –0.01 | –0.02 |
| 021 | 0.04 | –0.02 | 0 | 0.1 | –0.02 | –0.01 | –0.02 | 0 |
| 022 | 0.07 | 0.02 | 0.01 | 0.13 | 0 | –0.02 | –0.01 | –0.02 |
| 023 | 0.06 | 0 | 0 | 0.12 | 0.01 | –0.01 | –0.01 | –0.03 |
| 024 | 0.06 | –0.02 | 0.01 | 0.14 | –0.01 | 0 | –0.03 | –0.02 |
| 025 | 0.1 | –0.01 | 0 | 0.15 | –0.01 | –0.01 | 0 | –0.03 |
| 026 | 0.03 | –0.02 | 0 | 0.14 | –0.01 | –0.03 | –0.02 | 0.02 |
| 027 | 0.02 | –0.02 | 0 | 0.17 | –0.02 | –0.04 | –0.03 | 0.01 |
| 028 | 0.03 | –0.01 | 0 | 0.03 | 0.01 | 0 | 0 | –0.01 |
| 029 | 0.11 | –0.03 | 0 | –0.02 | –0.04 | 0.13 | 0.02 | –0.03 |
| 030 | 0.11 | –0.04 | 0.03 | –0.01 | –0.05 | 0.11 | 0.03 | –0.03 |
| 031 | 0.13 | –0.02 | 0.01 | 0.01 | –0.04 | 0.06 | 0.06 | –0.03 |
| 032 | 0.13 | –0.03 | 0.01 | –0.01 | –0.02 | 0.08 | 0.05 | –0.03 |
| 033 | 0.07 | –0.02 | 0.02 | –0.01 | –0.02 | 0.03 | 0.02 | 0.02 |
| 034 | 0.09 | 0 | 0 | 0 | 0 | 0.01 | 0.04 | 0.02 |
| 035 | 0.07 | 0 | 0.07 | –0.06 | –0.04 | 0.06 | –0.02 | 0 |
| 036 | 0.08 | 0.02 | 0.05 | –0.05 | –0.04 | 0.11 | –0.02 | 0.01 |
| 037 | 0.12 | –0.02 | –0.03 | –0.03 | –0.04 | –0.04 | 0.19 | –0.01 |
| 038 | 0.12 | –0.02 | –0.05 | –0.03 | –0.03 | –0.05 | 0.2 | –0.02 |
| 039 | 0.12 | –0.03 | –0.06 | –0.02 | –0.02 | –0.03 | 0.18 | –0.02 |
| 040 | 0.13 | –0.02 | –0.04 | –0.04 | –0.03 | –0.03 | 0.17 | 0 |
| 041 | 0.12 | –0.04 | –0.03 | –0.03 | –0.03 | –0.02 | 0.18 | –0.02 |
| 042 | 0.11 | –0.04 | –0.04 | –0.01 | –0.02 | –0.04 | 0.18 | –0.01 |
| 043 | 0.13 | –0.04 | –0.01 | 0 | –0.05 | 0.02 | 0.14 | –0.02 |
| 044 | 0.1 | –0.02 | –0.03 | 0.03 | 0.01 | 0.09 | –0.05 | 0.06 |
| 045 | 0.1 | –0.01 | –0.01 | 0.02 | 0.02 | 0.02 | –0.03 | 0.06 |
| 046 | 0.08 | 0 | 0 | 0 | –0.04 | 0.02 | –0.02 | 0.11 |
| 047 | 0.12 | –0.02 | –0.01 | 0.01 | –0.02 | 0.03 | 0.02 | 0.07 |
| 048 | 0.12 | –0.02 | –0.01 | –0.01 | –0.02 | 0.06 | 0.01 | 0.06 |
| 049 | 0.1 | –0.01 | –0.01 | 0.02 | 0.02 | 0.05 | –0.03 | 0.06 |
| 050 | 0.07 | 0.01 | –0.05 | –0.03 | –0.04 | 0.01 | 0.02 | 0.14 |
| 051 | 0.09 | –0.02 | 0.01 | 0.04 | –0.01 | 0.01 | 0 | 0.08 |
| 052 | 0.08 | 0.01 | –0.02 | 0.01 | –0.03 | –0.04 | –0.04 | 0.16 |
| 053 | 0.07 | 0.02 | –0.03 | 0 | –0.04 | –0.04 | –0.02 | 0.17 |
| 054 | 0.1 | 0 | –0.01 | 0 | –0.03 | –0.05 | 0 | 0.13 |
| 055 | 0.08 | 0.01 | –0.02 | 0.03 | –0.01 | –0.05 | –0.03 | 0.12 |
| 056 | 0.06 | 0.02 | –0.02 | 0 | –0.01 | –0.04 | –0.05 | 0.11 |
| 057 | 0.08 | 0.02 | –0.05 | –0.03 | –0.02 | –0.03 | 0 | 0.19 |
| 058 | 0.08 | 0.02 | –0.03 | 0.02 | –0.03 | –0.05 | –0.01 | 0.17 |
| 059 | 0.07 | 0.01 | –0.01 | –0.02 | –0.03 | –0.01 | 0.03 | 0.09 |
| 060 | 0.08 | –0.01 | 0 | –0.01 | –0.03 | –0.01 | 0.02 | 0.11 |
| 061 | 0.11 | 0.04 | 0.02 | 0.06 | –0.03 | –0.02 | 0.04 | –0.02 |
| 062 | 0.11 | 0.05 | 0.01 | 0.07 | –0.02 | –0.02 | 0.04 | –0.04 |
| 063 | 0.09 | 0.06 | 0.01 | 0.07 | 0 | –0.01 | 0.02 | –0.01 |
| 064 | 0.09 | 0.04 | 0.01 | 0.09 | 0 | –0.03 | 0.02 | –0.02 |
| 065 | 0.12 | 0.04 | 0.03 | 0.05 | –0.02 | –0.01 | 0.04 | –0.02 |

| | pF | ADHD | ANX | COND | PSYCH | DEP | MANIA | OCD |
|---|---|---|---|---|---|---|---|---|
| 066 | 0.08 | –0.02 | 0.04 | –0.02 | –0.04 | 0.08 | –0.01 | 0.02 |
| 067 | 0.1 | –0.03 | 0.05 | –0.02 | –0.02 | 0.05 | 0.01 | 0.02 |
| 068 | 0.09 | –0.02 | 0 | –0.01 | –0.01 | 0.06 | 0 | 0.02 |
| 069 | 0.05 | –0.01 | 0.09 | –0.01 | 0.01 | –0.04 | –0.01 | –0.02 |
| 070 | 0.06 | –0.02 | 0.1 | 0.01 | 0.01 | –0.04 | –0.02 | –0.01 |
| 071 | 0.06 | –0.01 | 0.06 | 0 | 0.01 | –0.03 | 0 | –0.01 |
| 072 | 0.05 | 0.01 | 0.1 | –0.01 | 0.02 | –0.04 | –0.04 | 0 |
| 073 | 0.04 | 0.02 | 0.07 | –0.03 | 0.01 | –0.03 | –0.01 | –0.01 |
| 074 | 0.07 | –0.01 | 0.08 | –0.01 | 0.01 | –0.03 | –0.03 | 0.02 |
| 075 | 0.07 | –0.01 | 0.1 | 0.02 | 0.01 | –0.04 | –0.03 | –0.01 |
| 076 | 0.07 | 0.02 | 0.06 | –0.03 | 0.01 | –0.01 | –0.03 | 0.03 |
| 077 | 0.1 | –0.01 | –0.01 | –0.02 | 0.09 | –0.02 | 0 | 0.01 |
| 078 | 0.09 | –0.03 | 0.01 | –0.01 | 0.05 | –0.02 | 0.05 | –0.01 |
| 079 | 0.1 | –0.01 | –0.01 | 0 | 0.07 | –0.01 | 0 | 0.01 |
| 080 | 0.07 | –0.02 | –0.01 | 0 | 0.03 | –0.02 | 0.04 | 0 |
| 081 | 0.11 | –0.03 | 0.01 | –0.01 | 0.06 | –0.03 | 0.05 | 0 |
| 082 | 0.09 | –0.03 | 0 | 0 | 0.01 | –0.01 | 0.03 | 0.02 |
| 083 | 0.03 | 0 | 0.01 | 0.01 | 0.01 | –0.01 | 0 | 0 |
| 084 | 0.08 | –0.03 | –0.01 | 0.06 | –0.01 | 0.04 | –0.01 | –0.01 |
| 085 | 0.06 | –0.03 | 0 | 0.11 | –0.02 | 0.02 | 0 | –0.03 |
| 086 | 0.05 | –0.02 | 0.02 | 0.05 | –0.03 | 0.06 | –0.03 | 0 |
| 087 | 0.05 | –0.03 | –0.02 | 0.12 | –0.01 | 0 | 0 | –0.01 |
| 088 | 0.03 | –0.03 | 0.01 | 0.05 | –0.02 | –0.01 | 0.02 | 0 |
| 089 | 0.07 | –0.04 | 0 | 0.08 | –0.01 | –0.01 | 0.02 | 0 |
| 090 | 0.07 | –0.04 | 0.02 | 0.05 | –0.01 | –0.02 | 0.04 | –0.02 |
| 091 | 0.08 | 0.04 | 0.01 | 0 | –0.01 | 0.08 | –0.02 | –0.02 |
| 092 | 0.06 | 0.08 | –0.02 | –0.04 | 0 | 0.06 | –0.01 | 0 |
| 093 | 0.05 | 0 | 0 | 0.01 | –0.01 | 0.09 | –0.05 | –0.02 |
| 094 | 0.07 | 0 | 0.06 | –0.01 | –0.02 | 0.02 | –0.01 | –0.02 |
| 095 | 0.06 | 0.01 | 0.08 | –0.01 | –0.03 | 0.02 | –0.01 | –0.01 |
| 096 | 0.06 | 0.02 | 0.09 | 0 | –0.02 | 0.02 | –0.02 | 0 |
| 097 | 0.09 | –0.01 | 0.06 | –0.01 | –0.02 | 0.01 | 0 | –0.02 |
| 098 | 0.04 | 0.03 | 0.07 | –0.03 | –0.02 | 0.02 | –0.01 | 0.01 |
| 099 | 0.11 | 0.16 | –0.01 | –0.03 | 0.01 | 0 | –0.02 | 0.01 |
| 100 | 0.13 | –0.02 | –0.01 | –0.02 | 0.11 | 0.02 | 0 | –0.03 |
| 101 | 0.08 | 0 | –0.01 | –0.01 | 0.13 | –0.03 | –0.04 | –0.01 |
| 102 | 0.11 | –0.01 | –0.02 | 0 | 0.13 | 0.01 | –0.05 | –0.02 |
| 103 | 0.11 | –0.02 | –0.02 | 0 | 0.12 | –0.03 | –0.02 | 0 |
| 104 | 0.13 | –0.02 | 0 | –0.02 | 0.13 | –0.01 | –0.02 | –0.03 |
| 105 | 0.09 | –0.01 | –0.02 | –0.01 | 0.13 | –0.04 | –0.03 | 0 |
| 106 | 0.11 | –0.03 | –0.01 | 0.02 | 0.1 | 0 | –0.04 | –0.01 |
| 107 | 0.09 | 0 | –0.01 | –0.01 | 0.15 | –0.05 | –0.03 | –0.02 |
| 108 | 0.13 | –0.02 | –0.03 | –0.02 | 0.13 | 0.02 | –0.02 | –0.01 |
| 109 | 0.12 | –0.01 | –0.02 | –0.03 | 0.14 | –0.03 | 0 | 0 |
| 110 | 0.11 | –0.02 | –0.01 | 0.01 | 0.11 | –0.03 | –0.02 | –0.01 |
| 111 | 0.11 | –0.01 | –0.04 | –0.03 | 0.1 | 0.06 | –0.05 | 0.02 |
| 112 | 0.08 | 0.01 | 0 | –0.02 | 0.02 | –0.01 | 0.08 | –0.01 |
| 113 | 0.08 | 0.02 | –0.01 | –0.02 | 0.02 | 0 | 0.07 | –0.01 |
| 114 | 0.08 | 0.04 | –0.01 | –0.02 | 0.05 | 0.01 | 0.03 | –0.01 |
| 115 | 0.09 | –0.02 | –0.04 | –0.03 | –0.01 | 0.14 | 0 | 0 |
| 116 | 0.09 | –0.02 | –0.03 | 0.03 | 0 | 0.09 | 0.02 | 0.01 |
| 117 | 0.11 | –0.01 | –0.05 | 0.01 | 0.01 | 0.15 | –0.01 | 0.01 |
| 118 | 0.07 | 0.02 | 0 | 0 | 0.05 | 0.07 | –0.05 | –0.02 |
| 119 | 0.06 | 0.04 | –0.02 | –0.05 | 0.01 | 0.09 | –0.05 | –0.03 |
| 120 | 0.06 | 0.03 | –0.03 | –0.03 | 0 | 0.07 | –0.04 | –0.02 |
| 121 | 0.08 | 0.05 | –0.03 | –0.03 | 0.01 | 0.11 | –0.07 | –0.02 |
| 122 | 0.09 | 0.05 | –0.02 | 0 | 0.03 | 0.11 | –0.05 | –0.01 |
| 123 | 0.08 | 0 | 0.13 | –0.01 | –0.01 | 0.03 | 0 | –0.04 |
| 124 | 0.07 | –0.01 | 0.11 | –0.02 | –0.01 | 0.02 | 0 | –0.04 |
| 125 | 0.07 | –0.02 | 0.13 | 0 | –0.01 | 0 | 0.02 | –0.05 |
| 126 | 0.09 | –0.02 | 0.14 | –0.01 | –0.01 | 0 | 0.01 | –0.05 |
| 127 | 0.1 | –0.02 | 0.13 | –0.01 | –0.01 | 0 | 0.02 | –0.05 |
| 128 | 0.1 | –0.02 | 0 | 0 | 0 | 0.07 | –0.02 | 0 |
| 129 | 0.09 | –0.02 | –0.01 | 0.01 | 0 | 0.13 | –0.04 | –0.02 |

**Supplementary Figure 3:** Principal component (pF) and independent component weights for each clinical item. pF: General psychopathology; ADHD: Attention Deficit Disorder; ANX: Anxiety; COND: Conduct disorder; PSYC: Psychosis; DEP: Depression; MANIA: Mania; OCD: Obsessive-compulsive disorder.

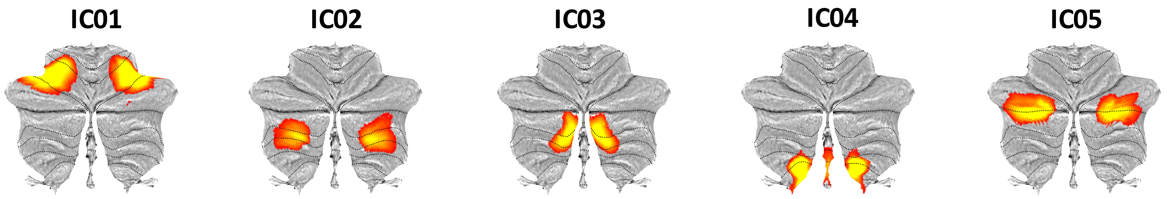


**Supplementary Figure 4:** ICA-decompositions of cerebellar grey matter maps using a model order of 5. Note IC01, which fuses regions assigned to cerebellar components IC03 and IC06 in the 10 component solution.

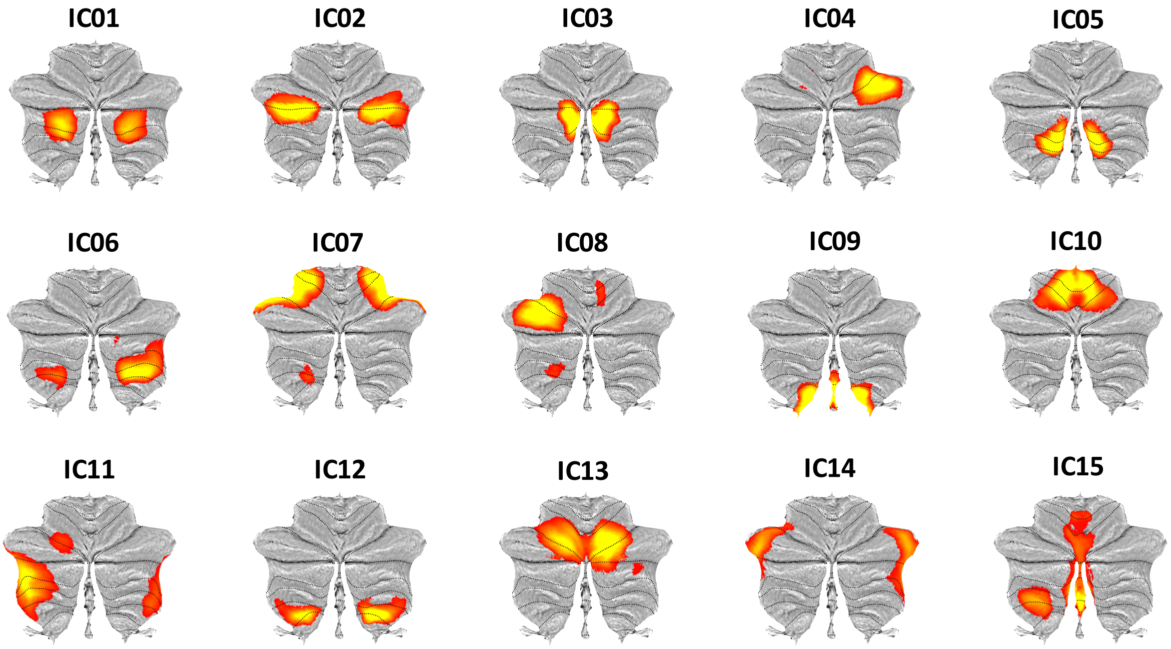


**Supplementary Figure 5:** ICA-decompositions of cerebellar grey matter maps using a model order of 15. Note IC04 and IC08, which largely correspond to the left and right aspect of the bilateral IC03 in the 10 component solution.

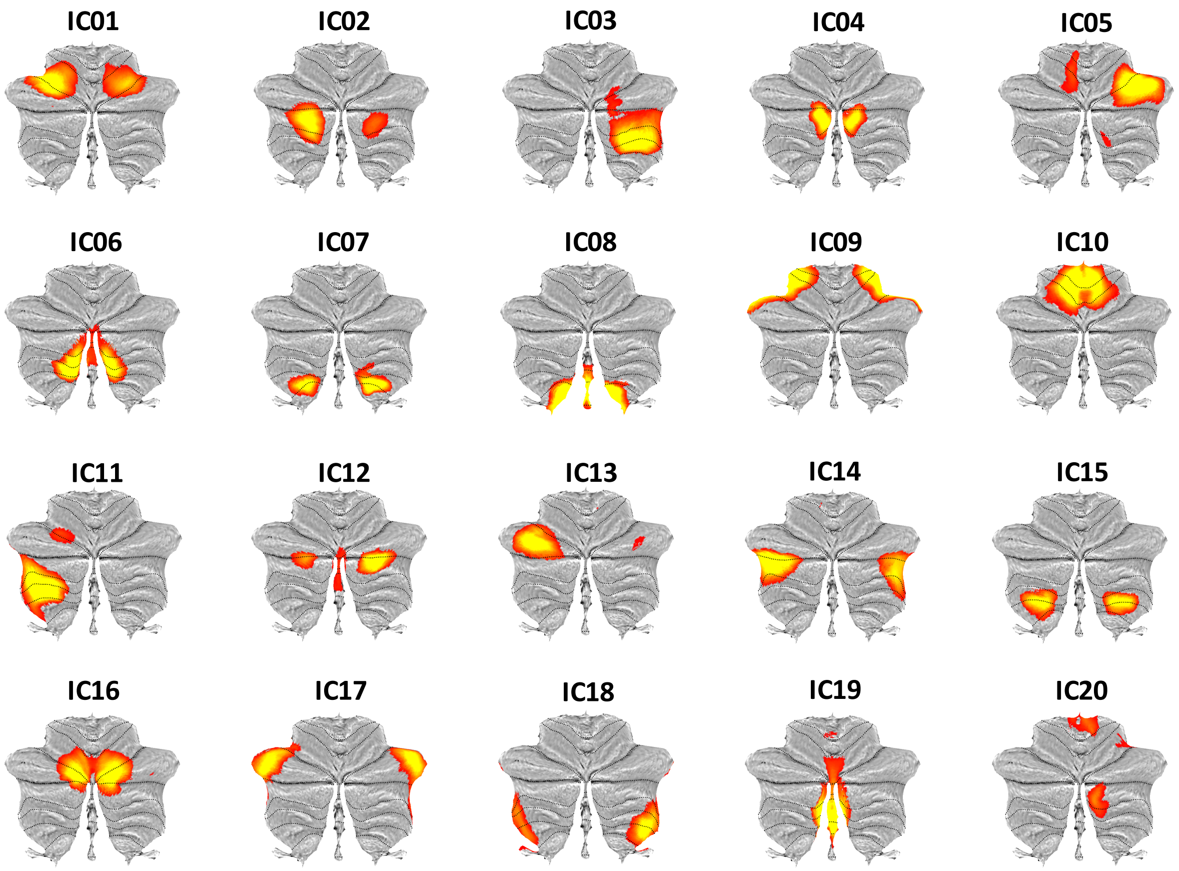


**Supplementary Figure 6**: ICA-decompositions of cerebellar grey matter maps using a model order of 20. Note IC05 and IC13, which largely correspond to the left and right aspect of the bilateral IC03 in the 10 component solution, and IC03 and IC11, which largely correspond to the left and right aspect of the bilateral IC03 in the 10 component solution.

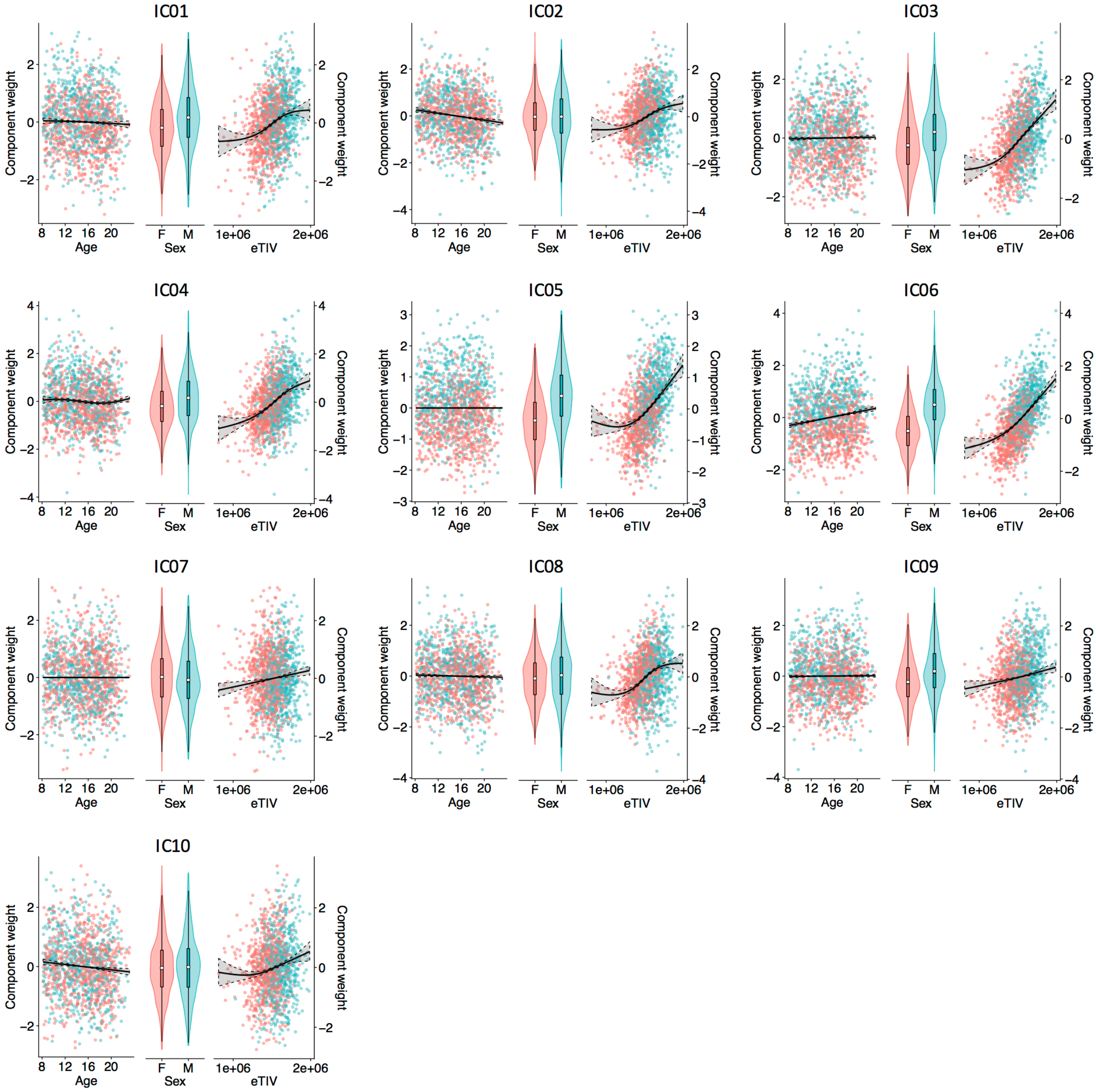


**Supplementary Figure 7:** Effects of age, gender and estimated total intracranial volume on subject weights for the 10 cerebellar components. Solid lines in the scatter plots depict the GAM-estimated smooth function, while shaded regions represent +/- 2 SEM. Distributions for each gender are represented in combined violin and box-plots, with the white dot indicating the group mean. For statistics, see Supplementary Table 4.

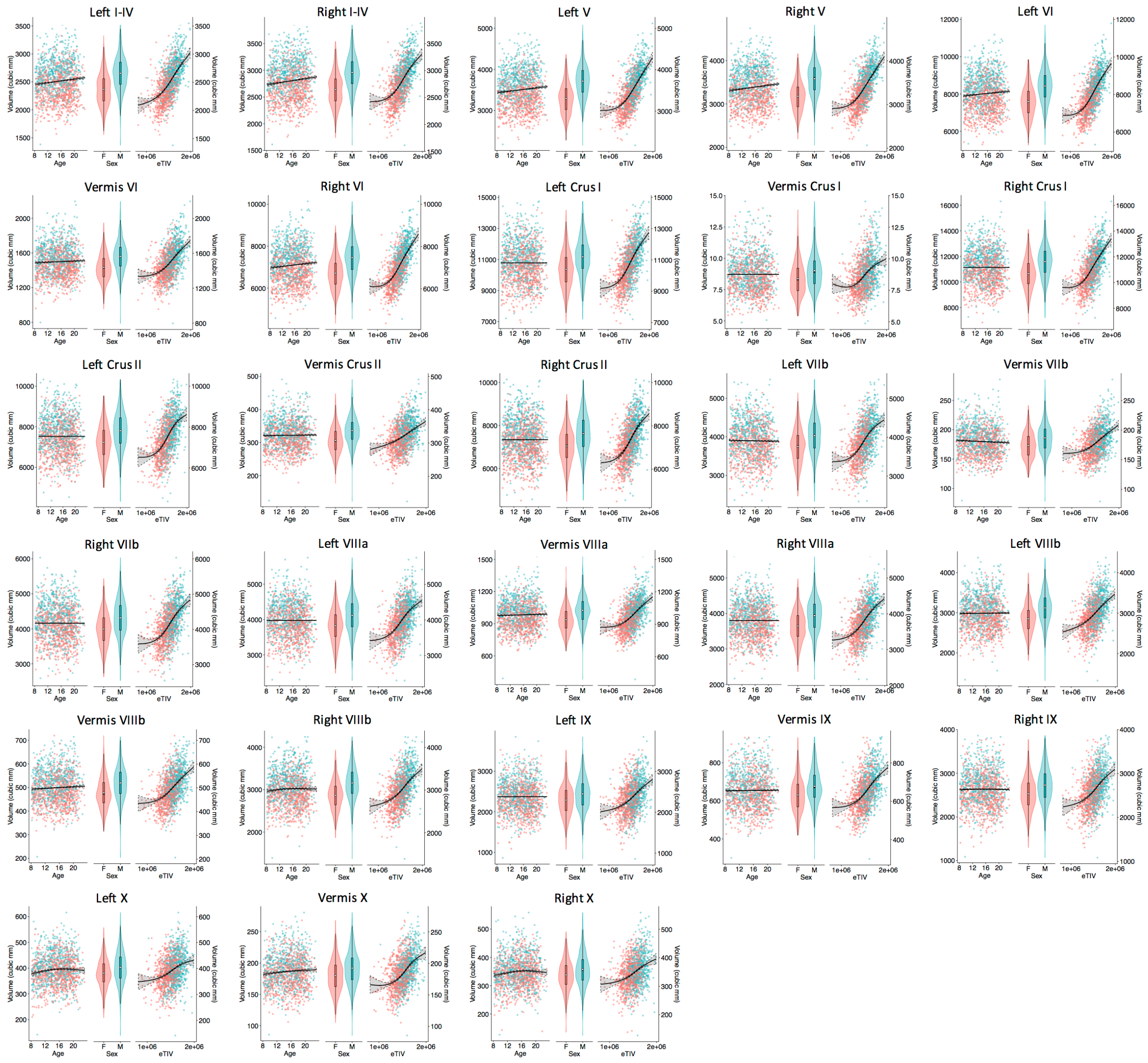


**Supplementary Figure 8:** Effects of age, gender and estimated total intracranial volume on volumes of the 28 cerebellar lobules. Solid lines in the scatter plots depict the GAM-estimated smooth function, while shaded regions represent +/- 2 SEM. Distributions for each gender are represented in combined violin and box-plots, with the white dot indicating the group mean. For statistics, see Supplementary Table 5.

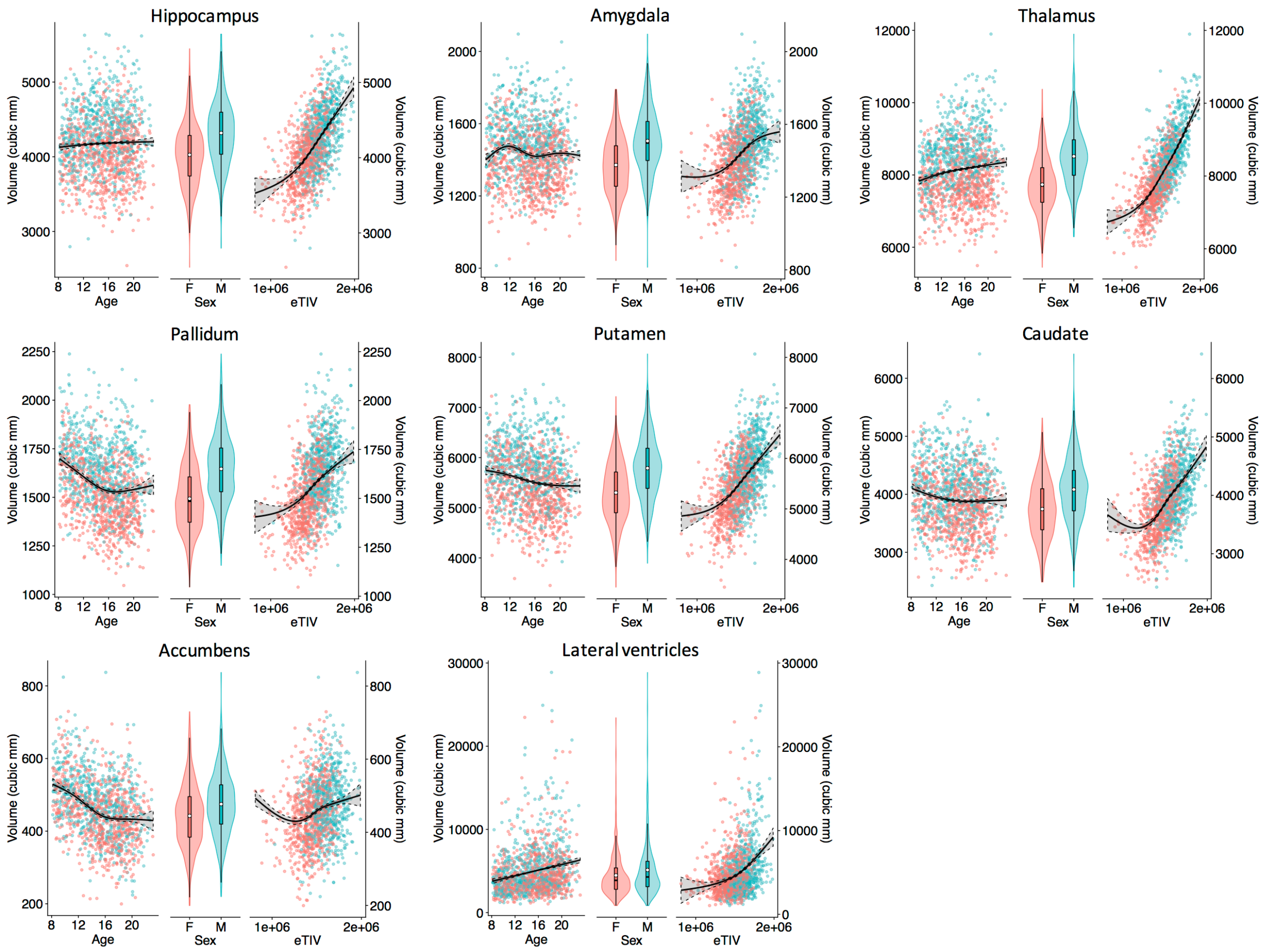


**Supplementary Figure 9:** Effects of age, gender and estimated total intracranial volume on volumes of the 8 (bilateral) subcortical structures. Solid lines in the scatter plots depict the GAM-estimated smooth function, while shaded regions represent +/- 2 SEM. Distributions for each gender are represented in combined violin and box-plots, with the white dot indicating the group mean. For statistics, see Supplementary Table 6.

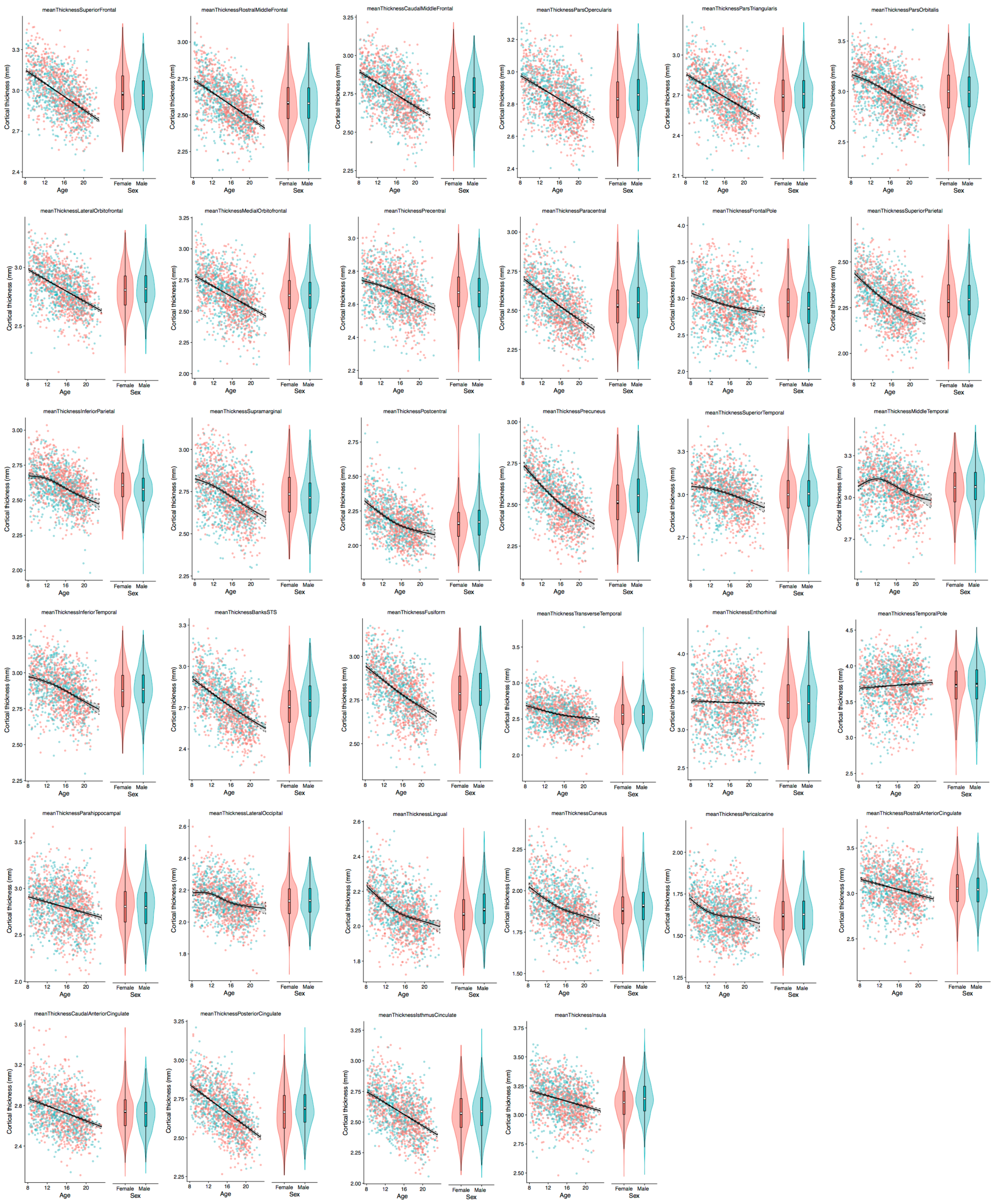


**Supplementary Figure 10:** Effects of age and gender on cortical thickness estimates for the 34 (bilateral) cerebro-cortical ROIs. Solid lines in the scatter plots depict the GAM-estimated smooth function, while shaded regions represent +/- 2 SEM. Distributions for each gender are represented in combined violin and box-plots, with the white dot indicating the group mean. For statistics, see Supplementary Table 7.

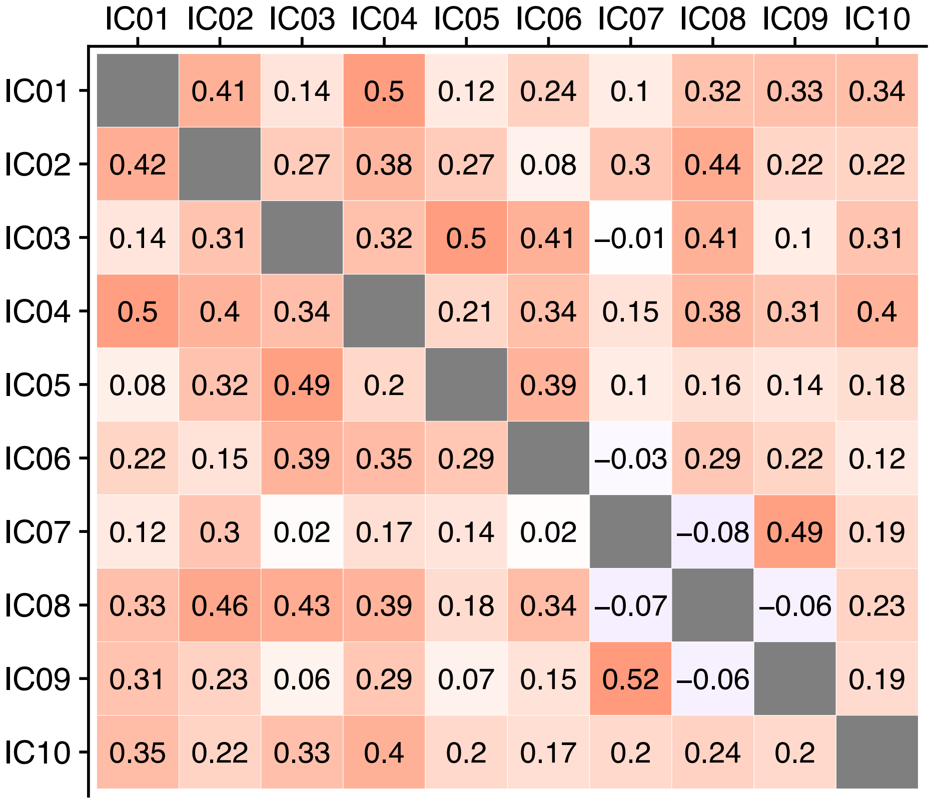


**Supplementary Figure 11:** Correlations between the 10 cerebellar components using raw components weights (above the diagonal) and component weights adjusted for effects of sex, age and total estimated intracranial volume (below the diagonal).

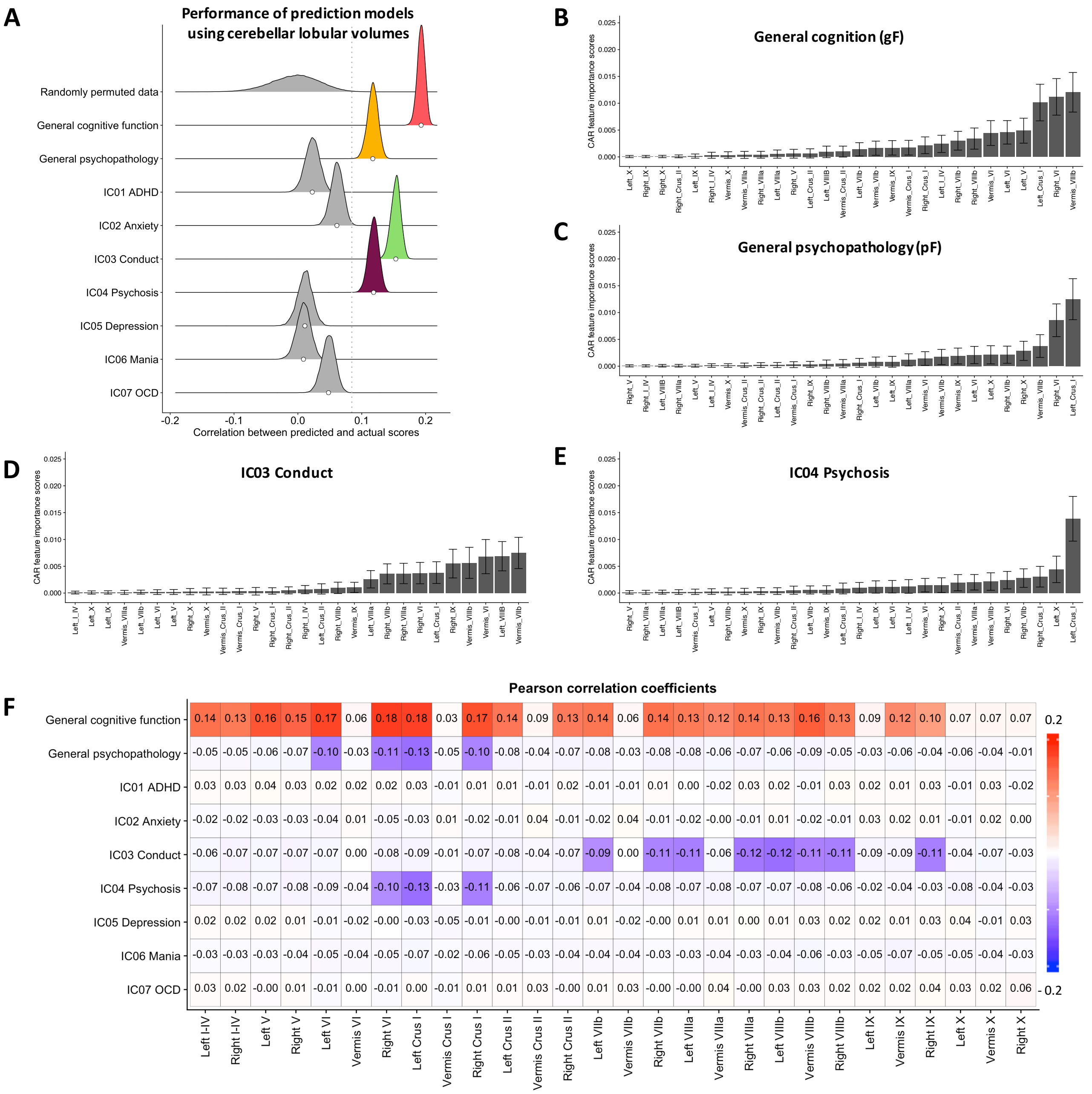


**Supplementary Figure 12:** Results from prediction models using cerebellar lobules **(A).** Feature importance weights (CAR-scores) for the four significant prediction models using the volumes of 28 cerebellar lobular ROIs **(B-E)**. CAR-scores were computed for each of 10,000 iterations of the model on randomly 10-fold partitioned data, yielding 100,000 estimates for each model. Error bars denote the 2.5th and 97.5th percentiles of these CAR-score distributions. Results from univariate correlation analyses **(F)**. Colored tiles mark significant associations ($p < .05$, based on 10,000 permutations and corrected for multiple comparisons across the matrix).

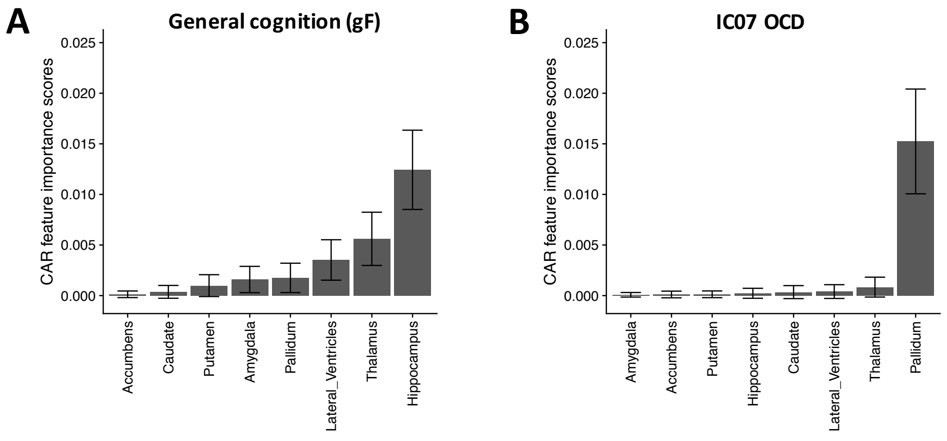


**Supplementary Figure 13:** Feature importance weights (CAR-scores) for the two significant prediction models using subcortical volumes. CAR-scores were computed for each of 10,000 iterations of the model on randomly 10-fold partitioned data, yielding 100,000 estimates for each model. Error bars denote the 2.5th and 97.5th percentiles of these CAR-score distributions.

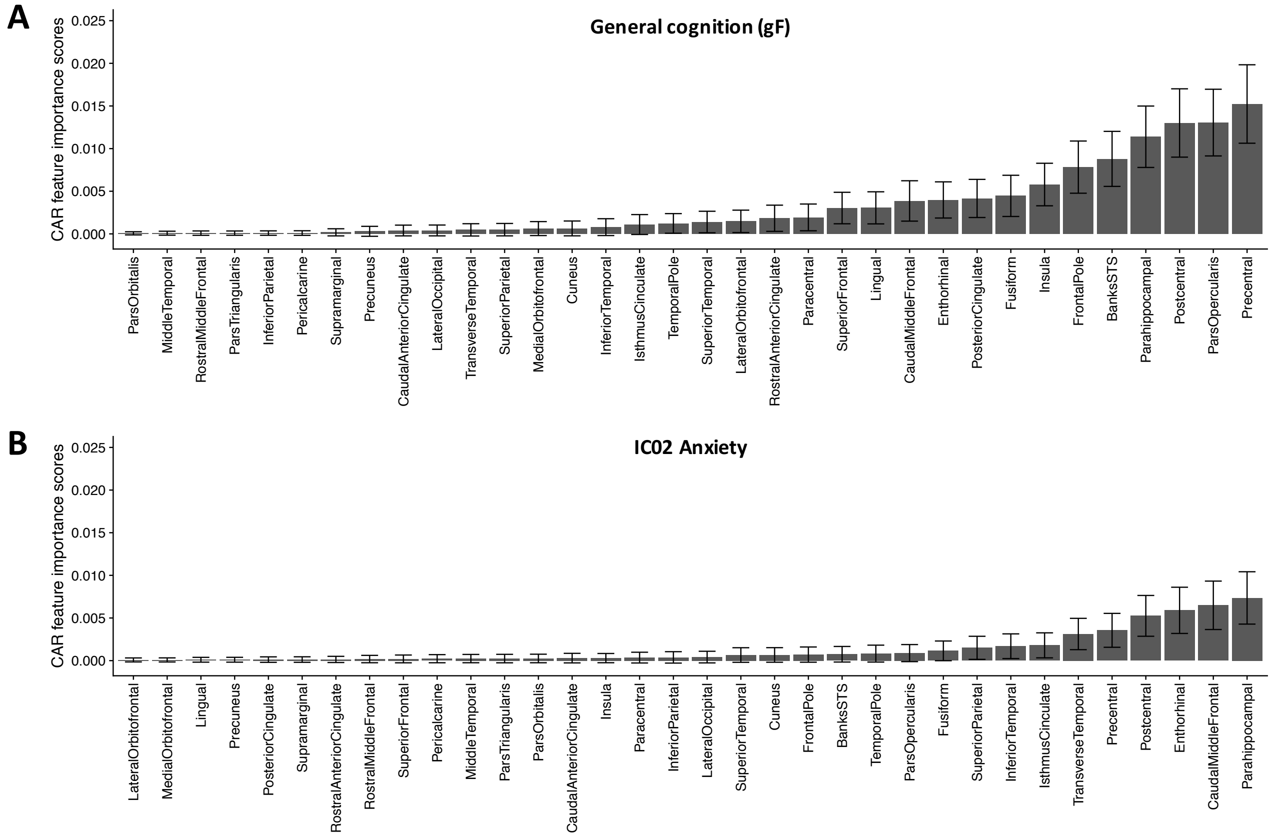


**Supplementary Figure 14:** Feature importance weights (CAR-scores) for the two significant prediction models using mean cortical thickness from 34 bilateral ROIs. CAR-scores were computed for each of 10,000 iterations of the model on randomly 10-fold partitioned data, yielding 100,000 estimates for each model. Error bars denote the 2.5th and 97.5th percentiles of these CAR-score distributions.

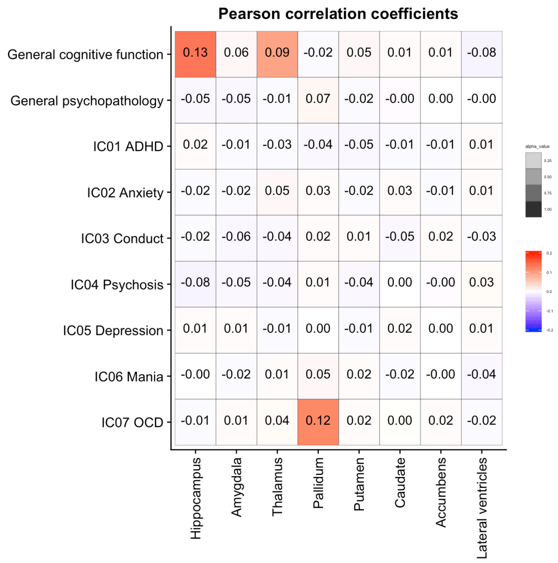


**Supplementary Figure 15:** Results from univariate correlation analyses using subcortical volumes. Colored tiles mark significant associations ($p$ < .05, based on 10,000 permutations and corrected for multiple comparisons across the matrix)

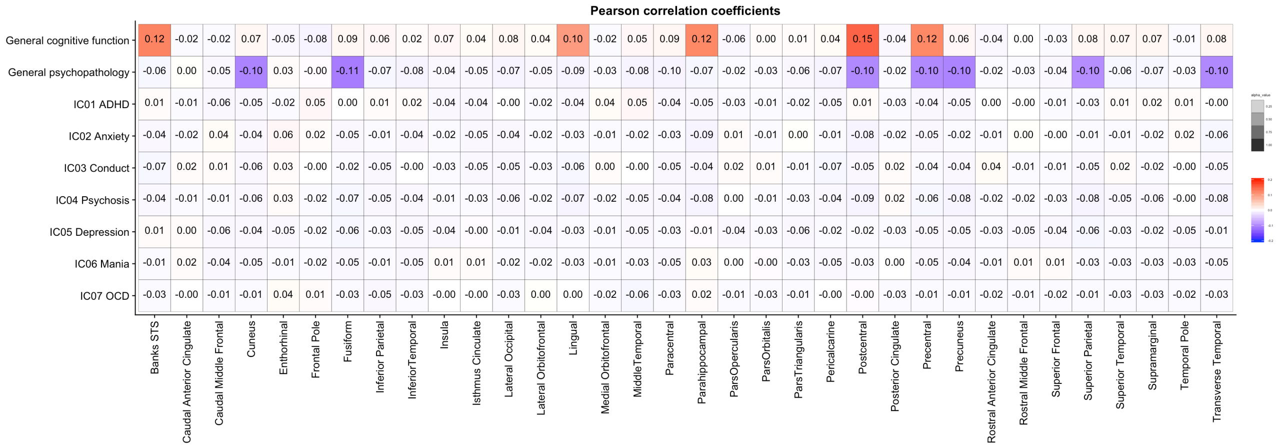


**Supplementary Figure 16:** Results from univariate correlation analyses using mean cortical thickness from 34 bilateral ROIs. Colored tiles mark significant associations ($p$ < .05, based on 10,000 permutations and corrected for multiple comparisons across the matrix)

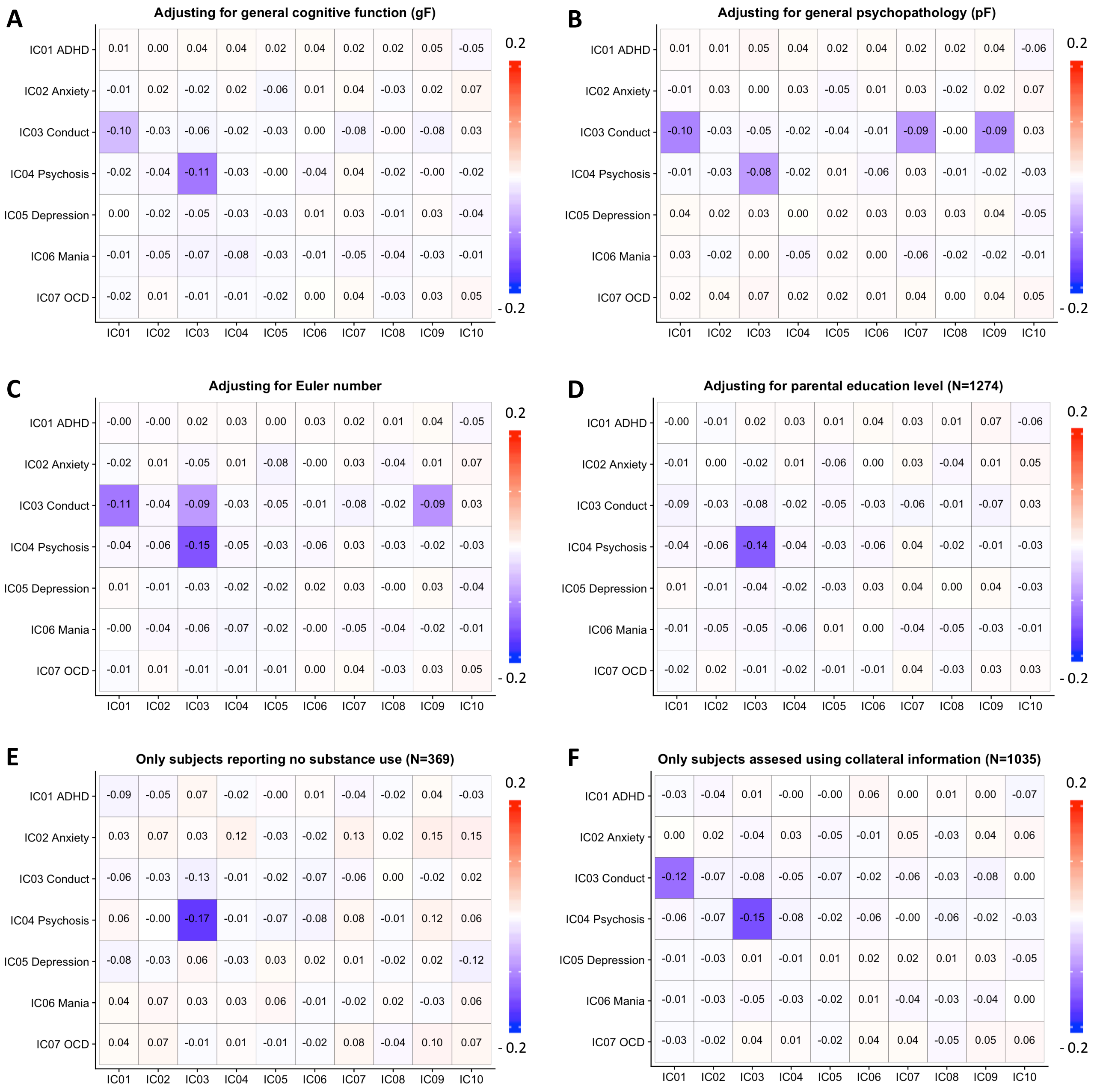


**Supplementary Figure 17:** Results from univariate control analyses using cerebellar independent components. Colored tiles mark significant associations (p< .05, corrected for multiple comparisons across the matrix).

**Supplementary Table 1:** 17 cognitive test scores included in the principal component analysis (PCA) and their factor loadings on the first principal component.

| Cognitive test | Included outcome measure | gF-weight |
|---|---|---|
| The Penn Age Differentiation Test | Percent correct responses | 0.2845 |
| The Penn Face Memory Test | Total correct responses | 0.1953 |
| Penn Emotion Identification Test | Total Correct Responses for All Test Trials, by genus | 0.1593 |
| Penn Word Memory Test | Total Correct Responses for All Test Trials | 0.1367 |
| Penn Verbal Reasoning Test | Total Correct Responses for All Test Trials, by genus | 0.2770 |
| Penn Emotion Differentiation Test | Percent of Correct Responses for All Test Trials, by genus | 0.2344 |
| Penn Motor Praxis task | Median Response Time for Correct Mouse Click Responses | -0.1565 |
| Penn Matrix Reasoning Test | Percent of Correct Responses for All Test Trials, by genus | 0.2613 |
| Finger Tapping Test | Sum of Mean of Tap Responses for Dominant Hand Trials and Mean of Tap Responses of Non-Dominant Hand Trials | 0.2210 |
| Visual Object Learning Test | Total Correct Responses for All Test Trial | 0.1505 |
| Letter N-Back Test | Number of Correct Responses to for 1-Back and2-Back Trials | 0.2939 |
| Penn Conditional Exclusion Test | Number of Categories Achieved | 0.1683 |
| Penn Conditional Exclusion Test | Calculated Accuracy Measure | 0.1183 |
| Penn Continuous Performance Test | Total of Correct Responses to Number Trials (TP) and Letter Trials (TP) | 0.1873 |
| Penn Continuous Performance Test | Median Response Time for Correct Responses to Number Trials (TP) and Letter Trials (TP) | -0.1901 |
| Penn Line Orientation Test | Percent Correct Responses for All Test Trials, by genus | 0.5059 |
| Wide Range Assessment Test(Reading/IQ) | WRAT: Wide Range Assessment Test 4 Total Raw Score | 0.2917 |

**Supplementary Table 2:** The 129 clinical items included in PCA and ICA decompositions.

| # | Questionnaire/Item |
|---|---|
| | **Attention deficit Hyperactivity Disorder** |
| 1 | Did you often have trouble paying attention or keeping your mind on your school, work, chores, or other activities that you were doing? |
| 2 | Did you often have problems following instructions and often fail to finish school, work, or other things you meant to get done? |
| 3 | Did you often dislike, avoid, or put off school or homework (or any other activity requiring concentration)? |
| 4 | Did you often lose things you needed for school or projects at home (assignments or books) or make careless mistakes in school work or other activities? |
| 5 | Did you often have trouble making plans, doing things that had to be done in a certain kind of order, or that had a lot of different steps? |
| 6 | Did you often have people tell you that you did not seem to be listening when they spoke to you or that you were daydreaming? |
| 7 | Did you often have difficulty sitting still for more than a few minutes at a time, even after being asked to stay seated, or did you often fidget with your hands or feet or wiggle in your seat or were you "always on the go"? |
| 8 | Did you often blurt out answers to other people's questions before they finished speaking or interrupt people abruptly? |
| 9 | Did you often join other people's conversations or have trouble waiting your turn (e.g., waiting in line, waiting for a teacher to call on you in class)? |
| | **Agoraphobia** |
| 10 | Have you ever been very nervous or afraid of: being in crowds (for example, a classroom, cafeteria, restaurant, or movie theater)? |
| 11 | Have you ever been very nervous or afraid of: going to public places (such as a store or shopping mall)? |
| 12 | Have you ever been very nervous or afraid of: being in an open field? |
| 13 | Have you ever been very nervous or afraid of: going over bridges or through tunnels? |
| 14 | Have you ever been very nervous or afraid of: traveling by yourself? |
| 15 | Have you ever been very nervous or afraid of: traveling away from home? |
| 16 | Have you ever been very nervous or afraid of: traveling in a car? |
| 17 | Have you ever been very nervous or afraid of: using public transportation like a bus or SEPTA? |
| | **Conduct disorder** |
| 18 | Was there ever a time when you often did things that got you into trouble with adults like lying or stealing (something worth more than $5, from family, others, or stores)? |
| 19 | Did you ever skip school, stay out at night later than you were supposed to (more than 2 hours), or run away from home overnight? |
| 20 | Did you ever set fires, break into cars, or destroy someone else's property on purpose? |
| 21 | Do you have a probation officer or have you ever been on probation? |
| 22 | Did you often bully others (hitting, threatening or scaring someone who was younger or smaller), threaten or frighten someone on purpose, or often start physical fights with others? |
| 23 | Have you ever been physically cruel to an animal or person (on purpose)? |
| 24 | Did you ever: try to hurt someone with a weapon (a bat, brick, broken bottle, knife, or gun)? |
| 25 | Did you ever: threaten someone? |
| 26 | Conduct Disorder: Did you ever: hold someone up? |
| 27 | Conduct Disorder: Did you ever: attack someone to steal from them? |
| 28 | Did you ever: trick or threaten someone into having sex with you, or did anyone ever accuse you of making them do something sexual? |
| | **Depression** |
| 29 | Has there ever been a time when you felt sad or depressed most of the time? |
| 30 | Has there ever been a time when you cried a lot, or felt like crying? |
| 31 | Has there ever been a time when you felt grouchy, irritable or in a bad mood most of the time; even little things would make you mad? |
| 32 | Has there ever been a time when nothing was fun for you and you just weren't interested in anything? |
| | **Eating Disorder** |
| 33 | Was there ever a time when you felt really fat or heavy, but other people said that you were too thin? |
| 34 | Has there been a time when your eating was out of control - you'd eat a large amount of food in a short period of time and could not stop yourself? |
| | **Generalized Anxiety** |
| 35 | Have you ever been a worrier? |
| 36 | Did you worry a lot more than most children/people your age? |
| | **Mania/Hypomania** |

| | |
|---|---|
| 37 | Have there been times when you were much more active, excited or energetic than usual, had problems sitting still, or needed to move around a lot? |
| 38 | Has there ever been a time when you felt so full of energy that you couldn't stop doing things and didn't get tired? |
| 39 | Has there ever been a time when you felt like you hardly needed sleep? |
| 40 | Have there been times when you kept talking a lot, couldn't stop talking, talked faster than usual, had thoughts faster than usual, or had so many ideas in your head that you could hardly keep track of them? |
| 41 | Have you ever had a time when you felt much more happy or excited than you usually do when there was nothing special going on? |
| 42 | Have you ever had a time when you felt like you could do almost anything? |
| 43 | Has there ever been a time when you felt unusually grouchy, cranky, or irritable; when the smallest things would make you really mad? |
| | **Obsessive Compulsive Disorder** |
| 44 | Have you ever been bothered by thoughts that don't make sense to you, that come over and over again and won't go away, such as concern with harming others/self? |
| 45 | Have you ever been bothered by thoughts that don't make sense to you, that come over and over again and won't go away, such as pictures of violent things? |
| 46 | Have you ever been bothered by thoughts that don't make sense to you, that come over and over again and won't go away, such as thoughts about contamination/germs/illness? |
| 47 | Have you ever been bothered by thoughts that don't make sense to you, that come over and over again and won't go away, such as fear that you would do something/say something bad without intending to? |
| 48 | Have you ever been bothered by thoughts that don't make sense to you, that come over and over again and won't go away, such as feelings that bad things that happened were your fault? |
| 49 | Have you ever been bothered by thoughts that don't make sense to you, that come over and over again and won't go away, such as forbidden/bad thoughts? |
| 50 | Have you ever been bothered by thoughts that don't make sense to you, that come over and over again and won't go away, such as need for symmetry/exactness? |
| 51 | Have you ever been bothered by thoughts that don't make sense to you, that come over and over again and won't go away, such as religious thoughts? |
| 52 | Have you ever had to do something over and over again - that would have made you feel really nervous if you couldn't do it, like: cleaning or washing (for example, your hands, house)? |
| 53 | Have you ever had to do something over and over again - that would have made you feel really nervous if you couldn't do it, like: counting? |
| 54 | Have you ever had to do something over and over again - that would have made you feel really nervous if you couldn't do it, like: checking (for example, doors, locks, ovens)? |
| 55 | Have you ever had to do something over and over again - that would have made you feel really nervous if you couldn't do it, like: getting dressed over and over again? |
| 56 | Have you ever had to do something over and over again - that would have made you feel really nervous if you couldn't do it, like: going in and out a door over and over again? |
| 57 | Have you ever had to do something over and over again - that would have made you feel really nervous if you couldn't do it, like: ordering or arranging things? |
| 58 | Have you ever had to do something over and over again - that would have made you feel really nervous if you couldn't do it, like: doing things over and over again at bedtime, like arranging the pillows, sheets, or other things? |
| 59 | Have you ever saved up so many things that people complained or they got in the way |
| 60 | Do you feel the need to do things just right (like they have to be perfect)? |
| | **Oppositional Defiant Disorder** |
| 61 | Was there a time when you often did things that got you into trouble with adults such as losing your temper, arguing with or talking back to adults, or being grouchy or irritable with them? |
| 62 | Was there a time when you often got into trouble with adults for refusing to do what they told you to do or for breaking rules at home/school? |
| 63 | Did you often annoy other people on purpose or blame other people for your mistakes (excluding siblings)? |
| 64 | Did you ever get into trouble for getting even with other people by doing things to hurt them, telling lies about them, or messing up their things? |

| | |
|---|---|
| 65 | Were you often irritable or grouchy, or did you often get angry because you thought that things were unfair? |
| | **Panic Disorder** |
| 66 | Have you ever had an attack like this? |
| 67 | Has there ever been a time when all of a sudden you felt very, very scared or uncomfortable - and your chest hurt, you couldn't catch your breath, your heart beat very fast, you felt very shaky, and sweaty/tingly/numb in your hands or feet? |
| 68 | Has there ever been a time when all of a sudden, you felt that you were losing control, something terrible was going to happen, that you were going crazy, or going to die? |
| | **Specific phobia** |
| 69 | Have you ever been very nervous or afraid of animals or bugs, like dogs, snakes, or spiders? |
| 70 | Have you ever been very nervous or afraid of being in really high places, like a roof or tall building? |
| 71 | Have you ever been very nervous or afraid of water or situations involving water, such as a swimming pool, lake, or ocean? |
| 72 | Have you ever been very nervous or afraid of storms, thunder, or lightning? |
| 73 | Have you ever been very nervous or afraid of doctors, needles, or blood? |
| 74 | Have you ever been very nervous or afraid of closed spaces, like elevators or closets? |
| 75 | Have you ever been very nervous or afraid of flying or airplanes? |
| 76 | Have you ever been very nervous or afraid of any other things or situations? |
| | **Psychosis** |
| 77 | Have you ever heard voices when no one was there? |
| 78 | Has there ever been anything unusual about the way things smelled or felt or looked? |
| 79 | Have you ever seen visions or seen things which other people could not see? |
| 80 | Have you ever smelled strange odors other people could not smell? |
| 81 | Have you ever had strange feelings in your body like things were crawling on you or someone touching you and nothing or no one was there? |
| 82 | Have you ever believed in things and later found out they weren't true, like people being out to get you, or talking about you behind your back, or controlling what you do or think? |
| | **Post-traumatic Stress Disorder** |
| 83 | Have you ever been in a flood or a tornado or an earthquake or a hurricane or some other natural disaster where you thought you were going to die or be seriously hurt? |
| 84 | Have you ever been in a situation where you thought you or someone close to you was going to be killed or be hurt very badly (e.g. family violence)? |
| 85 | Have you ever been attacked by somebody or badly beaten? |
| 86 | Have you ever been very upset by someone forcing you to do something sexual? |
| 87 | Have you ever been threatened with a weapon? |
| 88 | Have you ever been in a bad accident? |
| 89 | Other than television or at the movies, have you ever seen or heard somebody get killed or get hurt very badly or die? |
| 90 | Have you ever been very upset by seeing a dead body or by seeing pictures of the dead body of somebody you knew well? |
| | **General Probes** |
| 91 | Have you ever talked to a counselor, psychologist, social worker, psychiatrist or some other professional about your feelings or problems with your mood or behaviors? |
| 92 | Are you currently taking medication because of your emotions and/or behaviors? |
| 93 | Have you ever had to go to a hospital and stay overnight because of problems with your mood, feelings, or how you were acting? |
| | **Separation Anxiety** |
| 94 | Since you were 5 years old, has there ever been a time when you had a lot of worries about your (attachment figures) and were very upset or got sick (for example, felt sick to your stomach, headaches, thrown-up) when you were away from him/her? |
| 95 | Has there ever been a time when you wanted to stay home from school or not go to other places (for example, sleep-overs) without your (attachment figures)? |
| 96 | When you knew that you were going to be away from home or (attachment figure(s)), did you get very upset and worry (e.g., when you learned (attachment figure(s)) were going on an upcoming trip or night out)? |
| 97 | Did you ever worry/have bad dreams about something terrible happening to you or your (attachment figures) so that you would not see them again? |
| 98 | Were you scared to be alone in your room (or any place in your house) or did you need your (attachment figure(s)) to stay with |

| | |
|---|---|
| | you while you fell asleep? |
| | **Structural Interview for Prodromal Symptoms (Upper case items denote assessment by clinicians)** |
| 99 | TROUBLE WITH FOCUS AND ATTENTION: Severity Scale |
| 100 | I think that I have felt that there are odd or unusual things going on that I can't explain. |
| 101 | I think that I might be able to predict the future. |
| 102 | I may have felt that there could possibly be something interrupting or controlling my thoughts, feelings, or actions. |
| 103 | I have had the experience of doing something differently because of my superstitions. |
| 104 | I think I may get confused at times whether something I experience or perceive may be real or may be just part of my imagination or dreams. |
| 105 | I have thought that it might be possible that other people can read my mind, or that I can read others' minds |
| 106 | I wonder if people may be planning to hurt me or even may be about to hurt me. |
| 107 | I believe that I have special natural or supernatural gifts beyond my talents and natural strengths. |
| 108 | I think I might feel like my mind is "playing tricks" on me. |
| 109 | I have had the experience of hearing faint or clear sounds of people or a person mumbling or talking when there is no one near me. |
| 110 | I think that I may hear my own thoughts being said out loud. |
| 111 | I have been concerned that I might be "going crazy." |
| 112 | Do people ever tell you that they can't understand you? |
| 113 | Do people ever seem to have difficulty understanding you? |
| 114 | CHANGES IN SPEECH, DISORGANIZED COMMUNICATION, TANGENTIAL SPEECH: Severity Scale |
| 115 | Do you ever feel a loss of sense of self or feel disconnected from yourself or your life? |
| 116 | Has anyone pointed out to you that you are less emotional or connected to people than you used to be? |
| 117 | CHANGES IN PERCEPTION OF SELF, OTHERS, OR THE WORLD IN GENERAL: Severity Scale |
| 118 | EXPRESSION OF EMOTION: Severity Scale |
| 119 | Within the past 6 months, are you having a harder time getting your work or schoolwork done? |
| 120 | Within the past 6 months, are you having a harder time getting normal activities done? |
| 121 | OCCUPATIONAL FUNCTIONING: Severity Scale |
| 122 | AVOLITION: Severity Scale |
| | **Social Phobia** |
| 123 | Was there ever a time in your life when you felt afraid or uncomfortable or really, really shy with people, like meeting new people, going to parties, or eating or drinking, writing or doing homework in front of others? |
| 124 | Was there ever a time in your life when you felt afraid or uncomfortable talking on the telephone or with people your own age who you don't know very well? |
| 125 | Was there ever a time in your life when you felt afraid or uncomfortable when you had to do something in front of a group of people, like speaking in class? |
| 126 | Was there ever a time in your life when you felt afraid or uncomfortable acting, performing, giving a talk/speech, playing a sport or doing a musical performance, or taking an important test or exam (even though you studied enough)? |
| 127 | Was there ever a time in your life when you felt afraid or uncomfortable because you were the center of attention and were concerned something embarrassing might happen and you felt very afraid or felt uncomfortable? |
| | **Suicidal Thoughts** |
| 128 | Have you ever thought a lot about death or dying? |
| 129 | Have you ever thought about killing yourself? |

**Supplementary Table 3:** Results from GAM-models of cognitive/clinical scores.

| Model | Adjusted $r^2$ | Deviance explained | Sex | | Age | |
|---|---|---|---|---|---|---|
| | | | t-value | p-value | F-value | p-value |
| **gF Cognition** | 0.409 | 41.1 % | ***4.019*** | ***6.15e-05*** | ***241*** | ***< 2e-16*** |
| **pF Psychopathology** | 0.0705 | 7.32 % | 0.297 | 0.767 | ***26.77*** | ***< 2e-16*** |
| **IC01 ADHD** | 0.0358 | 3.72 % | ***4.616*** | ***4.27e-06*** | ***6.994*** | ***< 8.56e-08*** |
| **IC02 Anxiety** | 0.0653 | 6.75 % | ***-8.910*** | ***< 2e-16*** | ***4.96*** | ***< 2.06e-05*** |
| **IC03 Conduct** | 0.059 | 6.17 % | ***4.497*** | ***7.48e-06*** | ***18.49*** | ***< 2e-16*** |
| **IC04 Psychosis** | 0.0604 | 6.35 % | ***2.481*** | ***0.0132*** | ***20.93*** | ***< 2e-16*** |
| **IC05 Depression** | 0.0781 | 8.09 % | -0.751 | 0.45302 | ***29.26*** | ***< 2e-16*** |
| **IC06 Mania** | 0.0339 | 3.64 % | ***2.489*** | ***0.0129*** | ***11.49*** | ***1.53e-11*** |
| **IC07 OCD** | 0.00667 | 0.797 % | ***-2.285*** | ***0.02243*** | ***1.127*** | ***0.0191*** |

**Supplementary Table 4:** Correlations between raw (upper triangle) and age and sex-adjusted (lower triangle) cognitive/clinical scores.

| | gF | pF | IC01 | IC02 | IC03 | IC04 | IC05 | IC06 | IC07 |
|---|---|---|---|---|---|---|---|---|---|
| **gF Cognition** | | 0.055 | -0.163 | -0.078 | 0.010 | -0.111 | 0.204 | 0.160 | 0.062 |
| **pF Psychopathology** | -0.147 | | 0.175 | 0.329 | 0.257 | 0.540 | 0.437 | 0.459 | 0.456 |
| **IC01 ADD** | -0.102 | 0.222 | | 0.011 | 0.01 | -0.034 | -0.003 | -0.038 | -0.030 |
| **IC02 Anxiety** | -0.142 | 0.329 | 0.049 | | -0.025 | 0.073 | -0.056 | -0.069 | 0.062 |
| **IC03 Conduct** | -0.146 | 0.238 | 0.022 | 0.003 | | -0.000 | 0.037 | -0.029 | 0.023 |
| **IC04 Psychosis** | -0.230 | 0.538 | -0.036 | 0.063 | 0.009 | | 0.066 | 0.004 | 0.094 |
| **IC05 Depression** | 0.080 | 0.418 | 0.039 | -0.066 | -0.024 | 0.088 | | 0.067 | 0.106 |
| **IC06 Mania** | 0.063 | 0.435 | -0.023 | -0.072 | -0.066 | -0.020 | 0.032 | | 0.057 |
| **IC07 OCD** | 0.033 | 0.459 | -0.013 | 0.046 | 0.019 | 0.097 | 0.092 | 0.052 | |

**Supplementary Table 5:** Results from GAM-models on cerebellar independent components.

| Model | Adjusted $r^2$ | Deviance explained | Sex | | Age | | eTIV | |
|---|---|---|---|---|---|---|---|---|
| | | | t-value | p-value | F-value | p-value | F-value | p-value |
| **IC01** | 0.089 | 9.21 % | 1.026 | 0.305 | 0.613 | 0.0812 | ***21.544*** | ***< 2e-16*** |
| **IC02** | 0.0759 | 7.91 % | ***-5.08*** | ***4.18e-07*** | ***6.674*** | ***1.57e-07*** | ***21.587*** | ***< 2e-16*** |
| **IC03** | 0.205 | 20.8 % | -0.538 | 0.590 | 0.186 | 0.186 | ***67.826*** | ***< 2e-16*** |
| **IC04** | 0.153 | 15.7 % | -1.648 | 0.0995 | ***1.823*** | ***0.0124*** | ***49.569*** | ***< 2e-16*** |
| **IC05** | 0.2772 | 27.4 % | ***6.739*** | ***2.33e-11*** | 0.002 | 0.317 | ***57.457*** | ***< 2e-16*** |
| **IC06** | 0.441 | 44.3 % | ***10.835*** | ***< 2e-16*** | ***15.33*** | ***5.25e-15*** | ***105,94*** | ***< 2e-16*** |
| **IC07** | 0.0103 | 1.16 % | ***-3.442*** | ***0.000595*** | 0 | 0.808544 | ***2.872*** | ***0.000424*** |
| **IC08** | 0.0985 | 10.4 % | ***-4.174*** | ***3.18e-05*** | 0.29 | 0.142 | ***36.49*** | ***< 2e-16*** |
| **IC09** | 0.057 | 5.87 % | ***4.792*** | ***1.83e-06*** | 0.076 | 0.253 | ***4.791*** | ***7.12e-06*** |
| **IC10** | 0.028 | 3.09 % | ***-2.569*** | ***0.0103*** | ***2.494*** | ***0.000913*** | ***7.814*** | ***1.47e-08*** |

**Supplementary Table 6:** Results from GAM-models on cerebellar lobules.

| Model | Adjusted $r^2$ | Deviance explained | Sex | | Age | | eTIV | |
|---|---|---|---|---|---|---|---|---|
| | | | t-value | p-value | F-value | p-value | F-value | p-value |
| **Left I to IV** | 0.436 | 43.8% | ***6.495*** | ***1.15e-10*** | ***5.114*** | ***3.89e-06*** | ***142.936*** | ***< 2e-16*** |
| **Right I to IV** | 0.445 | 44.7% | ***7.209*** | ***9.19e-13*** | ***6.425*** | ***2.7e-07*** | ***142.316*** | ***< 2e-16*** |
| **Left V** | 0.507 | 50.9% | ***7.847*** | ***8.42e-15*** | ***4.768*** | ***7.96e-06*** | ***187.116*** | ***< 2e-16*** |
| **Right V** | 0.507 | 50.8% | ***8.563*** | ***<2e-16*** | ***6.116*** | ***5.03e-07*** | ***179.262*** | ***< 2e-16*** |
| **Left VI** | 0.478 | 48% | ***4.937*** | ***8.87e-07*** | ***2.948*** | ***0.000363*** | ***188.797*** | ***< 2e-16*** |
| **Vermis VI** | 0.3 | 30.3% | ***3.853*** | ***0.000122*** | ***0.758*** | ***0.0449*** | ***86.209*** | ***< 2e-16*** |
| **Right VI** | 0.476 | 47.8% | ***4.86*** | ***1.31e-06*** | ***3.674*** | ***7.74e-05*** | ***188.113*** | ***< 2e-16*** |
| **Left Crus I** | 0.365 | 36.7% | 0.037 | 0.97 | 0.000 | 0.919 | ***144.9*** | ***< 2e-16*** |
| **Vermis Crus I** | 0.16 | 16.3% | 0.49 | 0.625 | 0.000 | 0.899 | ***46.61*** | ***< 2e-16*** |
| **Right Crus I** | 0.389 | 39.1% | 1.439 | 0.15 | 0.004 | 0.457 | ***150.879*** | ***< 2e-16*** |
| **Left Crus II** | 0.306 | 30.9% | 0.59 | 0.555 | 0.001 | 0.588 | ***108.442*** | ***< 2e-16*** |
| **Vermis Crus II** | 0.199 | 20.1% | ***5.774*** | ***9.55e-09*** | 0.158 | 0.203 | ***36.614*** | ***< 2e-16*** |
| **Right Crus II** | 0.323 | 32.5% | 0.825 | 0.41 | 0.0 | 0.659 | ***115.8*** | ***< 2e-16*** |
| **Left VIIb** | 0.272 | 27.4% | 0.454 | 0.65 | 0.137 | 0.219 | ***91.916*** | ***< 2e-16*** |
| **Vermis VIIb** | 0.192 | 19.5% | 1.739 | 0.0823 | ***0.855*** | ***0.0355*** | ***51.753*** | ***< 2e-16*** |
| **Right VIIb** | 0.298 | 30% | 0.724 | 0.469 | 0.003 | 0.464 | ***103.367*** | ***< 2e-16*** |
| **Left VIIIa** | 0.261 | 26.3% | 0.917 | 0.359 | 0.002 | 0.545 | ***84.861*** | ***< 2e-16*** |
| **Vermis VIIIa** | 0.266 | 26.8% | ***3.714*** | ***0.000212*** | 0.438 | 0.0976 | ***71.707*** | ***< 2e-16*** |
| **Right VIIIa** | 0.284 | 28.6% | 1.059 | 0.29 | 0.001 | 0.591 | ***94.416*** | ***< 2e-16*** |
| **Left VIIIb** | 0.252 | 25.5% | ***3.812*** | ***0.000144*** | 0.081 | 0.256 | ***65.826*** | ***< 2e-16*** |
| **Vermis VIIIb** | 0.251 | 25.4% | ***2.303*** | ***0.0214*** | ***1.061*** | ***0.0221*** | ***73.072*** | ***< 2e-16*** |
| **Right VIIIb** | 0.285 | 28.7% | ***4.992*** | ***6.73e-07*** | ***1.272*** | ***0.0299*** | ***71.249*** | ***< 2e-16*** |
| **Left IX** | 0.152 | 15.4% | -0.553 | 0.581 | 0.00 | 1 | ***47.44*** | ***< 2e-16*** |
| **Vermis IX** | 0.233 | 23.6% | -0.498 | 0.618 | 0.086 | 0.255 | ***79.439*** | ***< 2e-16*** |
| **Right IX** | 0.191 | 19.3% | 0.615 | 0.539 | 0.103 | 0.276 | ***57.121*** | ***< 2e-16*** |
| **Left X** | 0.116 | 12% | 0.961 | 0.337 | ***3.147*** | ***0.00049*** | ***28.428*** | ***< 2e-16*** |
| **Vermis X** | 0.214 | 21.7% | -0.812 | 0.417 | ***2.535*** | ***0.00102*** | ***70.884*** | ***< 2e-16*** |
| **Right X** | 0.12 | 12.4% | -0.019 | 0.984 | ***3.186*** | ***0.000427*** | ***32.475*** | ***< 2e-16*** |

**Supplementary Table 7:** Results from GAM-models on subcortical volumes.

| Model | Adjusted $r^2$ | Deviance explained | Sex | | Age | | eTIV | |
|---|---|---|---|---|---|---|---|---|
| | | | t-value | p-value | F-value | p-value | F-value | p-value |
| **Hippocampus** | 0.366 | 36.8 % | 0.957 | 0.339 | ***1.353*** | ***0.0154*** | ***139.471*** | ***< 2e-16*** |
| **Amygdala** | 0.26 | 26.3 % | ***6.464*** | ***1.41e-10*** | ***5.684*** | ***2.1e-05*** | ***48.137*** | ***< 2e-16*** |
| **Thalamus** | 0.592 | 59.4 % | ***4.596*** | ***4.7e-06*** | ***17.8*** | ***< 2e-16*** | ***309.2*** | ***< 2e-16*** |
| **Pallidum** | 0.351 | 35.4 % | ***8.112*** | ***1.08e-15*** | ***40.12*** | ***< 2e-16*** | ***54.17*** | ***< 2e-16*** |
| **Putamen** | 0.352 | 35.5 % | ***4.193*** | ***2.93e-05*** | ***13.43*** | ***1.31e-13*** | ***97.56*** | ***< 2e-16*** |
| **Caudate** | 0.323 | 32.7 % | 0.44 | 0.66 | ***6.678*** | ***3.62e-07*** | ***113.156*** | ***< 2e-16*** |
| **Accumbens** | 0.204 | 20.8 % | 1.858 | 0.0633 | ***58.84*** | ***< 2e-16*** | ***15.80*** | ***5.3e-15*** |
| **Lat. ventricles** | 0.155 | 15.7 % | ***-2.172*** | ***0.03*** | ***17.24*** | ***< 2e-16*** | ***40.27*** | ***< 2e-16*** |

**Supplementary Table 8:** Results from GAM-models of ROI mean cortical thickness.

| Model | Adjusted $r^2$ | Deviance explained | Sex | | Age | |
|---|---|---|---|---|---|---|
| | | | t-value | p-value | F-value | p-value |
| **Superior Frontal** | 0.29 | 29.1% | ***-3.97*** | ***7.56e-05*** | ***141.9*** | ***< 2e-16*** |
| **Rostral Middle Frontal** | 0.27 | 27.1% | ***-2.443*** | ***0.0147*** | ***129.7*** | ***< 2e-16*** |
| **Caudal Middle Frontal** | 0.22 | 21.9 % | -1.88 | 0.0604 | ***97.88*** | ***< 2e-16*** |
| **IFG Pars Opercularis** | 0.20 | 19.6 | ***2.089*** | ***0.0369*** | ***82.12*** | ***< 2e-16*** |
| **IFG Pars Triangularis** | 0.22 | 22.6 % | 0.328 | 0.743 | ***100.8*** | ***< 2e-16*** |
| **IFG Pars Orbitalis** | 0.18 | 18.4 % | -1.875 | 0.061 | ***78.29*** | ***< 2e-16*** |
| **Lateral Orbitofrontal** | 0.23 | 23.3 % | -0.115 | 0.908 | ***104.1*** | ***< 2e-16*** |
| **Medial Orbitofrontal** | 0.20 | 20 % | -1.191 | 0.234 | ***87.37*** | ***< 2e-16*** |
| **Precentral** | 0.10 | 10.4 % | -0.748 | 0.455 | ***40.08*** | ***< 2e-16*** |
| **Paracentral** | 0.28 | 28 % | 0.731 | 0.465 | **133.9** | ***< 2e-16*** |
| **Frontal Pole** | 0.06 | 6.57 % | ***-5.567*** | ***3.1e-08*** | ***17.88*** | ***< 2e-16*** |
| **Superior Parietal** | 0.26 | 26.1 % | -0.278 | 0.781 | ***122.2*** | ***< 2e-16*** |
| **Inferior Parietal** | 0.18 | 18.5 % | ***-5.906*** | ***4.41e-09*** | ***73.31*** | ***< 2e-16*** |
| **Supramarginal** | 0.17 | 17.3 % | ***-4.278*** | ***2.02e-05*** | ***70.23*** | ***< 2e-16*** |
| **Postcentral** | 0.22 | 22.1 % | -0.099 | 0.921 | ***97.91*** | ***< 2e-16*** |
| **Precuneus** | 0.36 | 36.6 % | ***3.698*** | ***0.000226*** | ***192.4*** | ***< 2e-16*** |
| **Superior Temporal** | 0.07 | 6.9 % | 0.194 | 0.846 | ***25.13*** | ***< 2e-16*** |
| **Middle Temporal** | 0.08 | 8.71 % | -0.369 | 0.712 | ***32.36*** | ***< 2e-16*** |
| **Inferior Temporal** | 0.15 | 14.8 % | 0.122 | 0.903 | ***59.66*** | ***< 2e-16*** |
| **Banks STS** | 0.29 | 29.3 % | ***3.607*** | ***0.000321*** | ***137.1*** | ***< 2e-16*** |
| **Fusiform** | 0.26 | 25.6 % | 1.712 | 0.0872 | ***117.2*** | ***< 2e-16*** |
| **Transverse Temporal** | 0.06 | 6.33 % | -0.513 | 0.608 | ***23.07*** | ***< 2e-16*** |
| **Entorhinal** | 0.001 | 0.23 % | -0.947 | 0.344 | 0.469 | 0.0902 |
| **Temporal Pole** | 0.004 | 0.52 % | 0.525 | 0.6 | ***1.589*** | ***0.00677*** |
| **Parahippocampal** | 0.05 | 5.17 % | -1.642 | 0.101 | ***18.56*** | ***< 2e-16*** |
| **Lateral Occipital** | 0.09 | 9.13 % | -0.383 | 0.702 | ***34.18*** | ***< 2e-16*** |
| **Lingual** | 0.21 | 20.9 % | ***3.066*** | ***0.00221*** | ***86.61*** | ***< 2e-16*** |
| **Cuneus** | 0.14 | 14.3 % | ***2.229*** | ***0.026*** | ***54.98*** | ***< 2e-16*** |
| **Pericalcarine** | 0.07 | 7.54 % | -0.282 | 0.778 | ***27.7*** | ***< 2e-16*** |
| **Rostral Anterior Cingulate** | 0.06 | 6.22 % | ***-2.021*** | ***0.0435*** | ***22.48*** | ***< 2e-16*** |
| **Caudal Anterior Cingulate** | 0.12 | 12.2 % | ***-3.235*** | ***0.00125*** | ***46.89*** | ***< 2e-16*** |
| **Posterior Cingulate** | 0.324 | 32.5 | **2.424** | **0.0155** | ***163.6*** | ***< 2e-16*** |
| **Isthmus Cingulate** | 0.23 | 23.1 % | 0.229 | 0.819 | ***104*** | ***< 2e-16*** |
| **Insula** | 0.08 | 8.11 % | ***3.29*** | ***0.00103*** | ***26.52*** | ***< 2e-16*** |

**Supplementary Table 9:** Significant results from the voxel-wise general linear models

| Contrast | Cluster extent (voxels) | Peak coordinates x | y | z | *t* | *p* | Anatomical labels |
|---|---|---|---|---|---|---|---|
| **gF positive** | 92614 | 36 | -41 | -28 | 8.29 | <.0001 | Widespread |
| | 41 | -20 | -32 | -38 | 5.62 | <.0001 | Left lobule X |
| **pF negative** | 5727 | 39 | -37 | -32 | 5.72 | <.0001 | Right Crus I, Right VI, Right Crus II |
| | 5220 | -42 | -71 | -24 | 6.83 | <.0001 | Left Crus I, Left Crus II, Left lobule VI |
| | 80 | -37 | -50 | -23 | 4.80 | .002 | Left lobule VI |
| | 64 | -23 | -74 | -20 | 4.59 | .006 | Left lobule VI |
| | 8 | -34 | -59 | -21 | 4.28 | .017 | Left lobule VI |
| | 7 | -27 | -72 | -44 | 4.30 | .016 | Left Crus II |
| | 3 | -32 | -64 | -58 | 4.24 | .021 | Left Lobule VIIb |
| **IC02 Anxiety negative** | 11 | -31 | -34 | -35 | 4.26 | .018 | Left lobule VI |
| **IC03 Conduct negative** | 6583 | 2 | -60 | -48 | 5.81 | <.0001 | Right Lobule VIII, Right Lobule IX, Right Lobule VIIb, Left Lobule IX, Vermis IX, Vermis VIIIa, Vermis VIIIb, Right Crus II |
| | 2732 | -26 | -49 | -57 | 5.40 | <.0001 | Left Lobule VIIIa, Left Lobule VIIb, Left Lobule IX |
| | 975 | -29 | -69 | -46 | 5.10 | .001 | Left Lobule VIIb, Left Crus II, Left VIIIa, Left Lobule VIIIb |
| | 950 | -43 | -61 | -37 | 4.93 | .001 | Left Crus I, Left Crus II |
| | 491 | -25 | -83 | -37 | 4.91 | .002 | Left Crus II |
| | 360 | -27 | -63 | -34 | 4.73 | .003 | Left Crus I, Left Lobule VI |
| | 193 | 36 | -40 | -28 | 4.57 | .006 | Right Lobule VI, Right Lobule V |
| | 38 | 32 | -82 | -25 | 4.29 | .016 | Right Crus I |
| | 31 | 20 | -87 | -35 | 4.30 | .016 | Right Crus II |
| | 23 | 33 | -76 | -31 | 4.27 | .018 | Right Crus I |
| | 19 | 28 | -63 | -35 | 4.25 | .019 | Right Crus I |
| | 14 | 38 | -76 | -26 | 4.22 | .021 | Right Crus I |
| | 11 | 4 | -40 | -18 | 4.47 | .009 | Right Lobule I to IV |
| | 9 | 32 | -68 | -20 | 4.31 | .016 | Right Lobule VI |
| | 3 | 21 | -57 | -29 | 4.21 | .021 | Right Lobule VI |
| **IC04 Psychosis negative** | 4182 | -39 | -71 | -26 | 5.63 | <.0001 | Left Crus I, Left Crus II, Left Lobule VI |
| | 3795 | 47 | -62 | -26 | 5.67 | <.0001 | Right Crus I, Right Lobule VI |
| | 39 | -42 | -41 | -35 | 4.99 | .002 | Left Crus I |
| | 1 | -22 | -74 | -21 | 4.22 | 0.021 | Left Lobule VI |
